# Cargo quantification of functionalized DNA origami for therapeutic application

**DOI:** 10.1101/2024.08.27.609963

**Authors:** Olivia J. Young, Hawa Dembele, Anjali Rajwar, Ick Chan Kwon, Ju Hee Ryu, William M. Shih, Yang C. Zeng

## Abstract

In recent years, notable advances in nanotechnology-based drug delivery have emerged. A particularly promising platform in this field is DNA origami-based nanoparticles, which offer highly programmable surfaces, providing precise control over the nanoscale spacing and stoichiometry of various cargo. These versatile particles are finding diverse applications ranging from basic molecular biology to diagnostics and therapeutics. This growing interest creates the need for effective methods to quantify cargo on DNA origami nanoparticles. Our study consolidates several previously validated methods focusing on gel-based and fluorescence-based techniques, including multiplexed quantification of protein, peptide, and nucleic acid cargo on these nanoparticles. This work may serve as a valuable resource for groups researchers keen on utilizing DNA origami-based nanoparticles in therapeutic applications.

**Graphical abstract:** 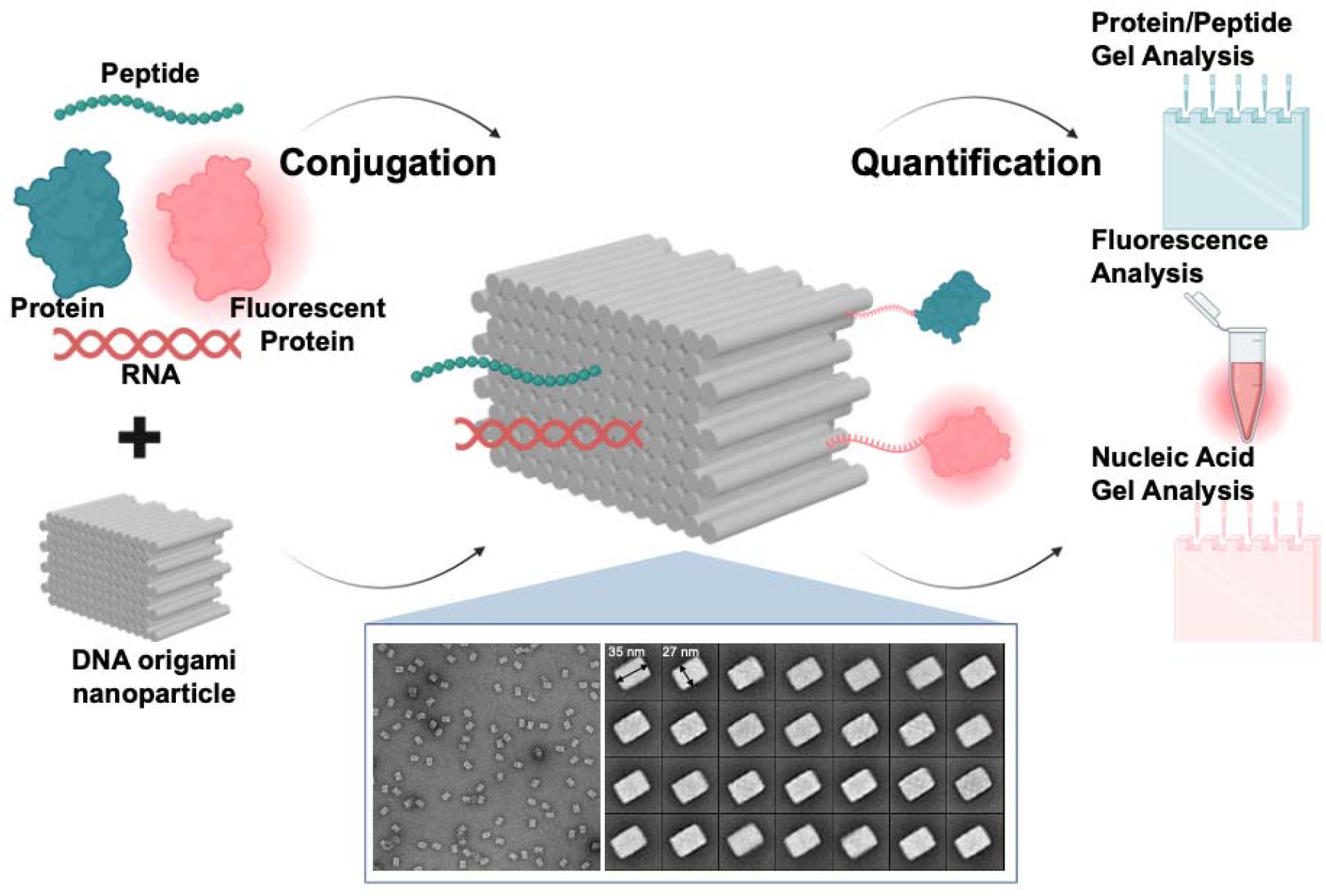

## Background

The potential of functional nanoparticles for drug delivery applications has attracted significant attention in recent years, partly due to their highly customizable nature. Various types of nanoparticles have been studied for their ability to carry bioactive cargos, making them suitable for a wide range of applications including therapeutics, diagnostics, and basic exploratory molecular biology^1-7^.Among these, DNA origami-based nanoparticles have emerged as a particularly promising platform. DNA origami utilizes a long single-stranded DNA ‘scaffold’ strand folded into a predetermined nanoscale configuration using short complementary oligonucleotide ‘staple strands’^8^. These nanoscale DNA origami structures can act as precise molecular pegboards, capable of organizing a wide array of cargos — from small molecules, nucleic acids, and peptides to whole proteins — with nanometer-scale precision. This highly customizable surface allows for the attachment of diverse cargos onto the self-assembled DNA origami nanoparticle^9-11^. To illustrate this, the DNA origami square block utilized here consists of 126 individual helices, providing more than 200 potential sites for facile conjugation^12,13^. These features make DNA origami a powerful and highly programmable technology for therapeutic delivery. Leveraging these capabilities, recent pre-clinical studies have highlighted DNA origami’s potential as a delivery platform for various therapeutic agents. The first cargo delivered to cells via DNA origami were intercalating chemotherapy drugs such as doxorubicin^14-17^, while other early studies focused on the delivery of nucleic acids such as CpG oligonucleotides^13,18-20^, aptamers^21-24^, and even DNA or siRNA for gene editing^25,26^. More recently, DNA origami has been used to deliver therapeutically relevant proteins and peptides^13,27-35^. Beyond direct therapeutic delivery, DNA origami has garnered interest for its ability to explore the spatial arrangement of antigens^36-38^ or adjuvant^13,39^ in vaccines, particularly for infectious diseases or cancers^13,18,38,40,41^. Additionally, DNA origami structures have been customized with complementary-shaped cavities for trapping viruses^42^. Research has also investigated the therapeutic potential of DNA origami in treating autoimmune disorders and thrombosis^24,43,44^.

DNA origami stands out in the degree of programmability in terms of nanoscale conjugation, setting it apart from other nanoparticle platforms^9-11^. While advanced techniques like DNA-PAINT and microscopy are invaluable for reporting nanoscale biology, DNA origami is complementary to these methods by enabling nanoscale perturbation of living systems, which is essential for understanding biological processes at the nanoscale^7,45^. Recent studies have leveraged the programmed nature of DNA origami to investigate how the spacing and quantity of cargo impact receptor binding efficiency and subsequent immune responses^7,46^. For instance, our group and others have demonstrated that nanometer-scale spacing of CpG on DNA origami nanoparticles significantly impacts receptor activation^39^ and anti-tumor immune responses in mice^13^. Similarly, Veneziano and colleagues demonstrated that placing five or more antigens on each origami particle, with an inter-antigen spacing of ∼25–30 □ nm, maximizes B cell activation and enhances the neutralizing antibody response^38,40^. Sun et al. demonstrated that incorporating 12 or more pMHC molecules per origami, along with anti-CD3 antibodies and co-stimulatory ligands on DNA origami-based artificial antigen-presenting cells, greatly increases T cell engagement and improves anti-cancer immune responses^29^. Other studies have demonstrated how the spatial arrangement of PD-1 and PD-L1 profoundly influences T cell function^6,47^, and how the spacing of DR5 and other tumor necrosis factor-related apoptosis-inducing ligands (TRAILs) influences cell apoptosis^46,48^.

These examples highlight the increasing interest in the use of DNA origami to study receptor engagement and downstream biological responses through the precise control of cargo spacing and copy number. To support the success of future research, especially studies focusing on DNA origami as a clinical therapeutic candidate, it is essential to have comprehensive methods for accurately confirming and characterizing cargo conjugation on DNA origami nanoparticles. Meeting Chemistry, Manufacturing, and Controls (CMC) standards for therapeutics necessitates exact product specifications, including precise quantification of therapeutic cargo, to ensure safety and consistency across production batches. Although fabrication and purification techniques for DNA origami have undergone extensive development and review^49-51^ in recent years, there remains a lack of studies compiling methodologies for characterizing conjugated cargo onto DNA origami particles^13,38,40^.

Here, we provide a practical resource highlighting gel-based and fluorescence-based methods for quantifying a wide array of cargo on DNA origami nanoparticles, including single-stranded DNA, double stranded RNA, peptides, and proteins. We demonstrate the simultaneous quantification of diverse cargos, emphasizing that multiplexed analytical capability will become increasingly important as DNA origami nanoparticle therapeutics evolve in complexity. This ensures a comprehensive assessment of the intricate cargo profiles associated with these nanoparticles.

## Results

### Conjugation and quantification of proteins on DNA origami nanoparticles

Previous studies from our group introduced the DNA origami square block (SQB), a square lattice DNA origami nanoparticle composed of 126 individual helices, offering 252 potential conjugation sites^12,13^. The SQB forms via self-assembly of a long ‘scaffold’ strand and multiple short oligonucleotide ‘staple’ strands. Specific staple strands are extended at either the 3’ or 5’ end to include ‘handle’ sequences to which a complementary ‘anti-handle’ strand bearing cargo can be attached via precise Watson and Crick base-pairing. Various cargos can be conjugated to the anti-handle strands through different chemical methods, enabling versatile attachment of different molecules to a single nanoparticle.

For our initial quantification experiment, we chose to conjugate a protein onto our SQB origami nanoparticle. Proteins are frequently used as cargo in therapeutic applications, serving roles as targeting agents, antigens, or immunostimulatory molecules. In this study, we used ovalbumin (OVA) as a model antigen. OVA was conjugated to an ‘anti-handle’ oligonucleotide through SMCC chemistry, and this conjugation was validated by SDS PAGE (Fig. 1A, B). The ‘anti-handle’ oligonucleotide-protein conjugate was then hybridized to the corresponding ‘handle’ strand on the DNA origami nanoparticle via Watson and Crick base pairing. Excess staple strands were removed using PEG precipitation (Fig. 1C). Successful hybridization was confirmed by observing a band shift on an agarose gel (Fig. 1D). Finally, DNase I was employed to degrade the DNA components of the protein-conjugated DNA origami nanoparticle, leaving behind only the conjugated protein, which was analyzed via SDS PAGE and silver stain (Fig. 1E). Using ImageJ analysis, we quantified an average of 11.8 proteins per SQB, despite being 24 available conjugation sites. We attribute this discrepancy primarily to steric hindrance caused by the large size of the OVA protein (Fig. 1F, Supplementary Table 1, Supplementary Fig. S1). We attempted to quantify the number of proteins on the SQB with OVA conjugated with Alexa Fluor 488 (OVA-AF488). However, the presence of AF488 interfered with the silver stain method and led to high background in the gel lane, limiting our ability to quantify the protein number (Supplementary Fig. 1).

**Figure 1.**
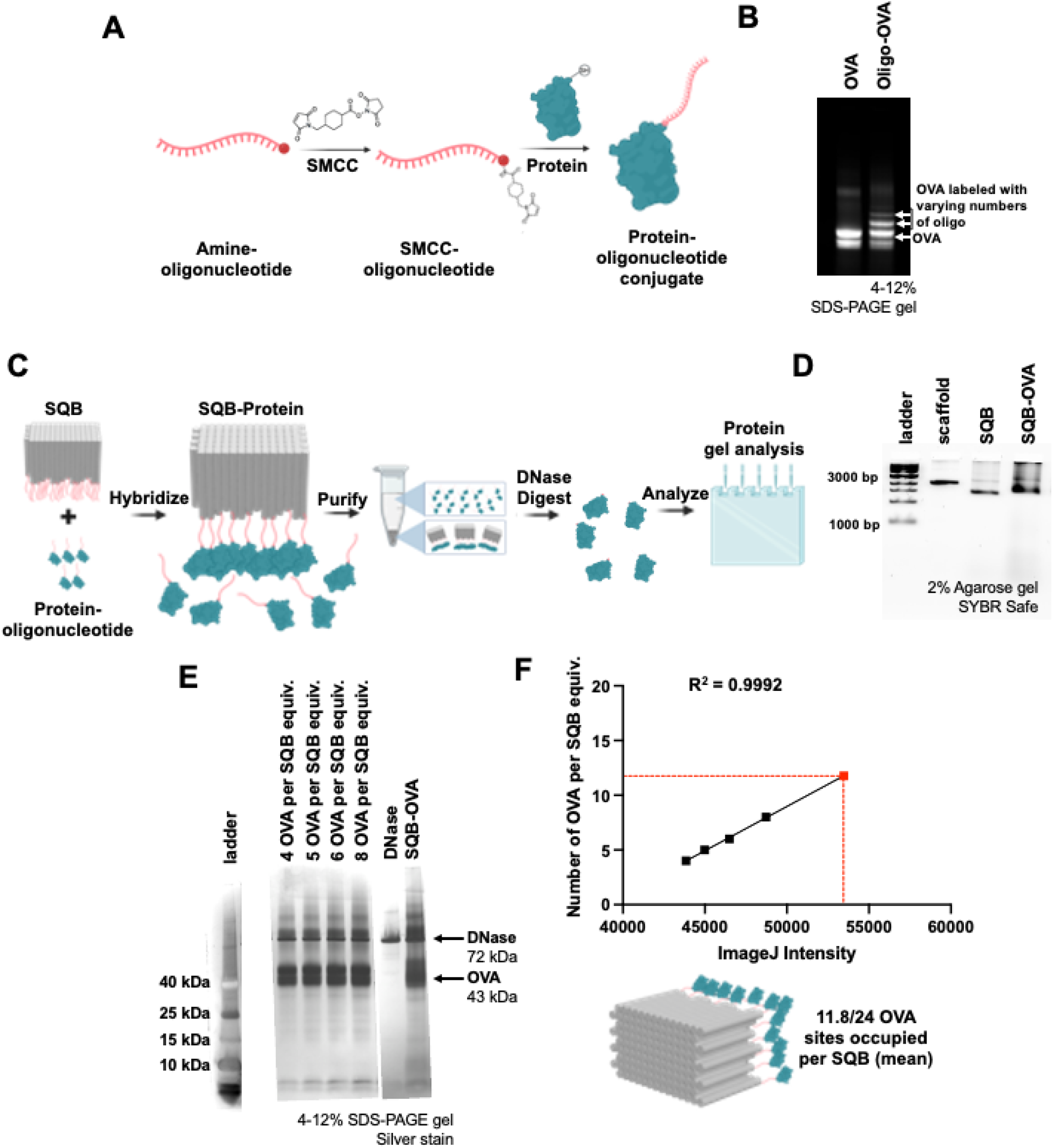
Quantifying protein cargo on DNA origami nanoparticles. (A) Schematic representation illustrating the oligonucleotide-protein conjugation reaction utilizing SMCC chemistry. (B) 15% dPAGE gel stained with SYBR Safe demonstrating successful protein-oligo conjugation. (C) Schematic representation showing the conjugation of the protein to the DNA origami SQB via Watson and Crick base pairing hybridization, followed by subsequent gel-based quantification analysis. (D) Agarose gel demonstrating the successful conjugation and purification of protein-bound SQBs. (E) Silver stain demonstrating successful conjugation and quantification of the OVA protein onto the SQB. The ladder, obtained from a different gel, is aligned using the DNase I band as the reference. The SQB-OVA and OVA dilution series were obtained from the same raw gel but were cropped to highlight the bands of interest; “equiv” refers to a unit of measure corresponding to the number of moles of SQB loaded in the rightmost lane. (F) Quantification of the OVA protein on the DNA origami SQB (4, 5, 6 and 8 OVA molecules per SQB) via linear regression of the ImageJ band intensity. The resulting relationship between the band intensity and the number of conjugated sites was used to determine the mean occupancy of the 24 designed OVA-capture sites on the DNA origami samples.

### Conjugation and quantification of fluorescent cargo on DNA origami nanoparticles

Building on our successful quantification of protein cargo, we aimed to demonstrate the ability to quantify fluorescently labeled protein cargo on DNA origami nanoparticles. To do this, OVA-AF488 protein was first conjugated to a DNA oligonucleotide using SMCC chemistry and then attached onto the SQB using handle/anti-handle hybridization (Fig. 2A-C). The resulting product was purified using PEG precipitation, and successful conjugation of OVA-AF488 protein was confirmed through a band shift on agarose gel (Fig. 2D). Excess OVA-AF488 protein was observed in the unpurified DNA origami SQB sample, as the fluorescence from AF488 overlaps with the wavelength region used to image SYBR-stained DNA. We performed a linear regression analysis comparing the AF488 fluorescence measured via Nanodrop to the known concentration of OVA-AF488 (Supplementary Table 2). This analysis allowed us to use the AF488 fluorescence of the OVA-conjugated SQB to quantify the number of OVA-AF488 molecules attached to the SQB origami nanoparticle (Fig. 2E). It’s important to note that this fluorescence analysis via Nanodrop is valid only within a specific dynamic range. When the measured fluorescence is too low, accurate quantification becomes difficult, as demonstrated by our dilution series measurements. Highly diluted samples showed vastly different values for the average number of OVA molecules per SQB compared to less diluted samples (Supplementary Table 3). The analysis showed an average of 12.9 OVA-AF488 molecules attached per SQB origami nanoparticle, which is consistent with the gel-based OVA quantification results described previously. The discrepancy between the number of conjugated OVA molecules (12.9) and the number of possible attachment sites (24) is likely due to steric hindrance. The large size of the OVA molecules prevents them from being closely packed on the extruding face of the SQB, as previously mentioned (Supplementary Tables 2–3).

**Figure 2.**
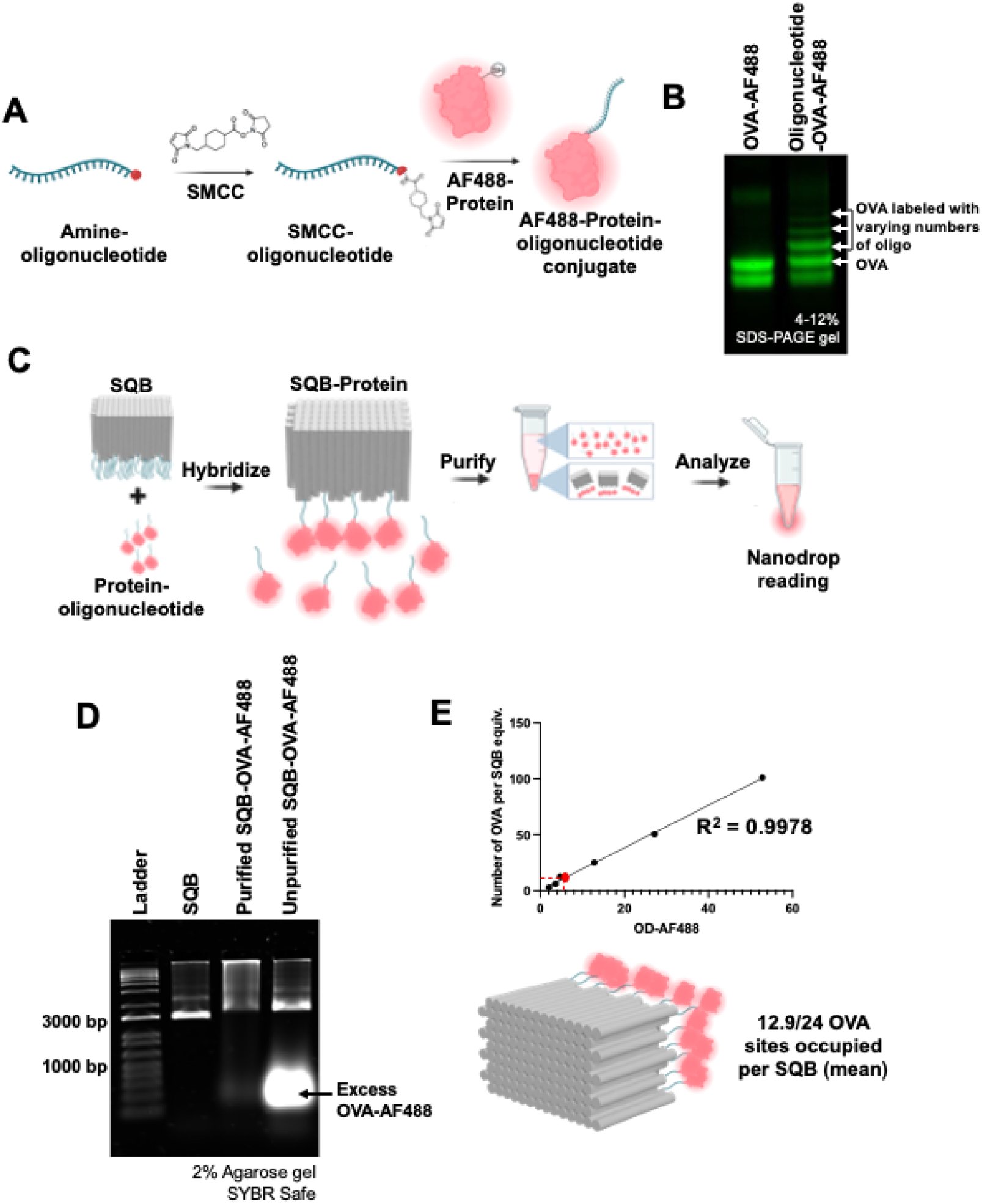
Quantifying fluorescent protein cargo on DNA origami nanoparticles. (A) Schematic representation illustrating the oligonucleotide-protein conjugation process utilizing the SMCC crosslinker. (B) 15% denaturing PAGE gel presenting the successful conjugation of fluorescent protein (OVA-AF488) to complementary anti-handle oligonucleotides. (C) Schematic representation showcasing the conjugation of protein onto the DNA origami SQB and the subsequent fluorescence-based quantification analysis. (D) Agarose gel demonstrating the successful conjugation and purification of fluorescent protein (OVA-AF488) oligonucleotides to the SQB nanoparticle. (E) Linear regression curve of OVA-AF488 fluorescence intensity used for quantifying the mean number of OVA molecules per SQB at different concentrations (0.275, 0.55, 1.1, 2.2, 4.4, and 8.8 µM).

### Conjugation and quantification of peptides on DNA origami nanoparticles

We successfully quantified protein conjugation onto DNA origami nanoparticles and aimed to demonstrate the parallel quantification of therapeutic peptides. We chose to use an immunostimulatory CD40 ligand cyclic peptide (CD40L) and quantify it alongside the previously discussed OVA protein antigen. Using the DBCO-azide click chemistry strategy, we conjugated the CD40L peptide to an ‘anti-handle’ oligonucleotide functionalized with an azide group (Fig. 3A)^52^. The successful conjugation and purification of the peptide to the oligonucleotide was confirmed via 15% denaturing PAGE (dPAGE) gel (Fig. 3B). These purified conjugates were then hybridized to the SQB nanoparticle. The successful hybridization was confirmed by observing a band shift on agarose gel (Fig. 3C, D). To quantify the cargo on the DNA origami nanoparticles, a DNase I digestion assay was performed as described in earlier sections. The resulting products were analyzed using SDS-PAGE, allowing for the effective multiplexed quantification of both the CD40L peptide and the OVA protein via ImageJ software (Fig. 3E, F). This method demonstrates simultaneous quantification of two types of therapeutic cargo on DNA origami nanoparticles. However, to obtain a detectable band intensity for the peptide, the protein sample tends to be oversaturated. This analysis estimated an average of 23.5 cyclic peptides and 11.8 OVA proteins conjugated to each SQB nanoparticle. The theoretical availability of only 18 sites for cyclic peptide conjugation suggests that the observed higher attachment might be due to the peptide’s non-specific adhesive properties. The quantified count of 11.8 OVA proteins per SQB is consistent with earlier data collected by gel and fluorescence quantification methods (Fig. 1 and 2), validating the reliability of these quantification methods (Supplementary Tables 4–5, Supplementary Figs. S2–S3).

**Figure 3.**
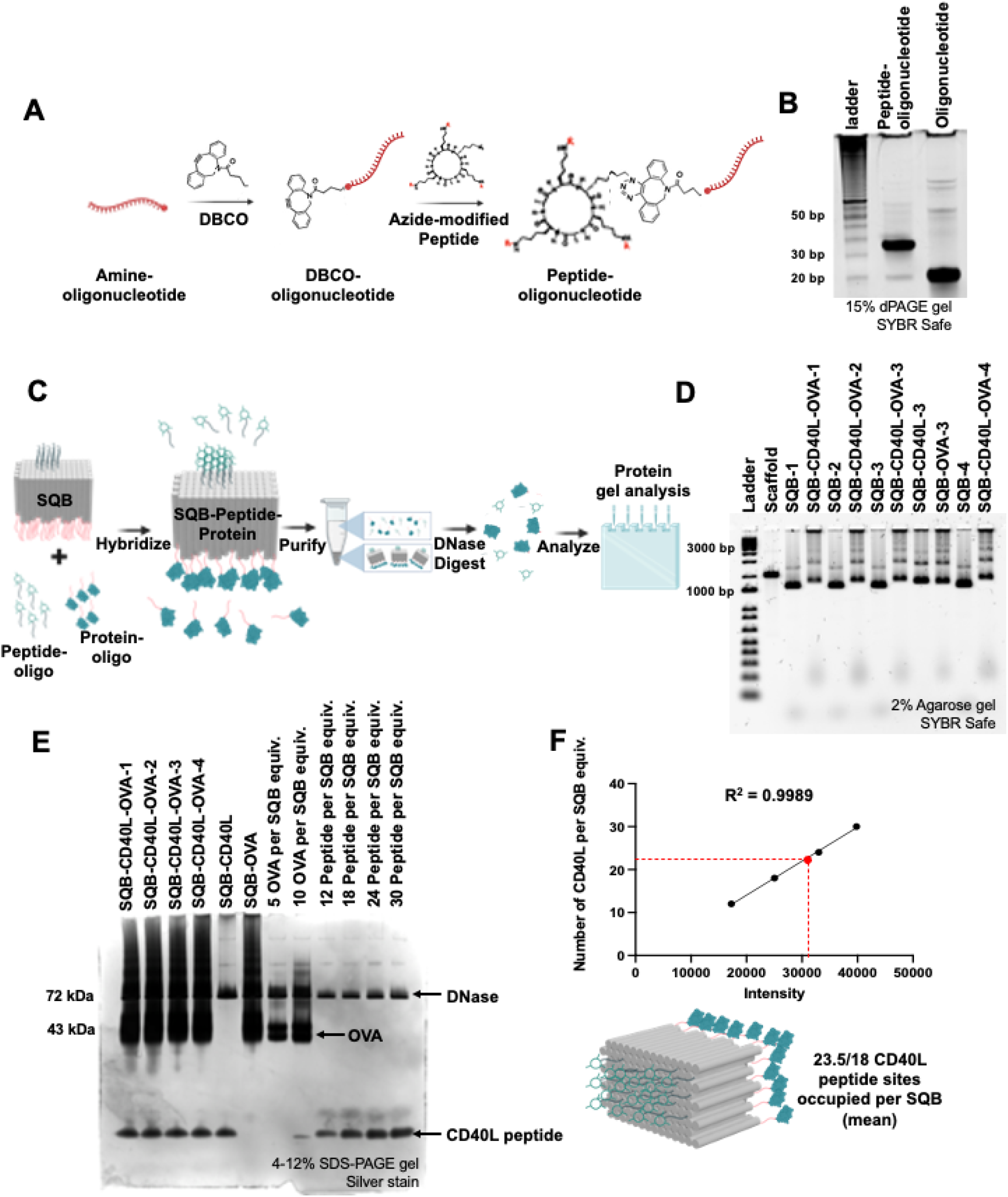
Quantifying peptide cargo on DNA origami nanoparticles. (A) Schematic representation illustrating the oligonucleotide-peptide conjugation process utilizing the dibenzocyclooctyne (DBCO)-Azide click chemistry. (B) 15% denaturing PAGE (dPAGE) gel stained with SYBR Safe, demonstrating successful peptide-oligo conjugation. (C) Schematic representation showcasing the conjugation of the peptide to the DNA origami SQB and the subsequent gel-based fluorescence analysis. (D) Agarose gel demonstrating the successful conjugation and purification of peptide-conjugated SQBs. SQB-CD40L-1 has extruding handles with 3.5 nm spacing; SQB-CD40L-2 has extruding handles with 5 nm spacing; SQB-CD40L-3 has extruding handles with 7 nm spacing; SQB-CD40L-4 has extruding handles with 10 nm spacing. SQB-1, SQB-2, SQB-3 and SQB-4 have the corresponding handles but are not conjugated with cargo. If only CD40L is delineated, then the SQB includes CD40L, but no conjugated OVA. If only OVA is delineated, then the SQB includes OVA, but no conjugated CD40L, even though the handles for CD40L conjugation are present. (E) Silver stain analysis demonstrating the successful conjugation and quantification of both the peptide and OVA protein onto the SQB. Here, “equiv” refers to a unit of measure corresponding to the number of moles of SQB loaded in the leftmost lane. (F) Quantification of the peptide on the DNA origami SQB using linear regression analysis of the ImageJ band intensity, measuring various concentrations (12, 18, 24, and 30 CD40L peptides per SQB). This analysis provided the relationship between the band intensity and the number of occupied peptide-capture sites, used to determine the occupancy of the 18 designed CD40L peptide-capture sites on the DNA origami samples.

### Conjugation and quantification of nucleic acids on DNA origami nanoparticles

To demonstrate the ability to conjugate and subsequently quantify double stranded RNA (dsRNA) molecules on the surface of the SQB origami, we designed two complementary single-stranded RNA (ssRNA) strands. One of these ssRNA strands contains an anti-handle DNA sequence that is complementary to a handle sequence on the SQB. Following duplexation, the RNA duplex was mixed with the SQB origami. The resulting dsRNA-SQB particles were purified, and the successful conjugation of the dsRNA was confirmed through a band shift on agarose gel (Fig. 4A–C). Following the procedures used for protein and peptide quantification, we performed a DNase digestion assay to reveal the dsRNA molecules that were originally attached to the surface of the SQB particles. In addition to the DNase-digested dsRNA-SQB sample, we prepared standard samples containing known amounts of dsRNA, corresponding to varying SQB conjugation equivalents, to serve as reference standards. The samples were then run on a denaturing PAGE gel and post-stained with SYBR Gold to visualize the dsRNA (Fig. 4D). The dsRNA band intensity was analyzed using ImageJ, and a linear regression analysis was applied to identify an equation describing the relationship between the number of conjugated dsRNA molecules and the band intensity (Fig. 4E). Of note, during the gel electrophoresis process, the denaturing conditions caused some of the RNA to denature, resulting in the appearance of three bands on the gel. Without denaturation, we would expect to observe only a single band corresponding to the RNA. Our analysis showed an average of 17.9 dsRNA molecules conjugated per SQB origami particle, out of a possible 18 conjugation sites (Supplementary Table 6, Supplementary Fig. S4).

**Figure 4.**
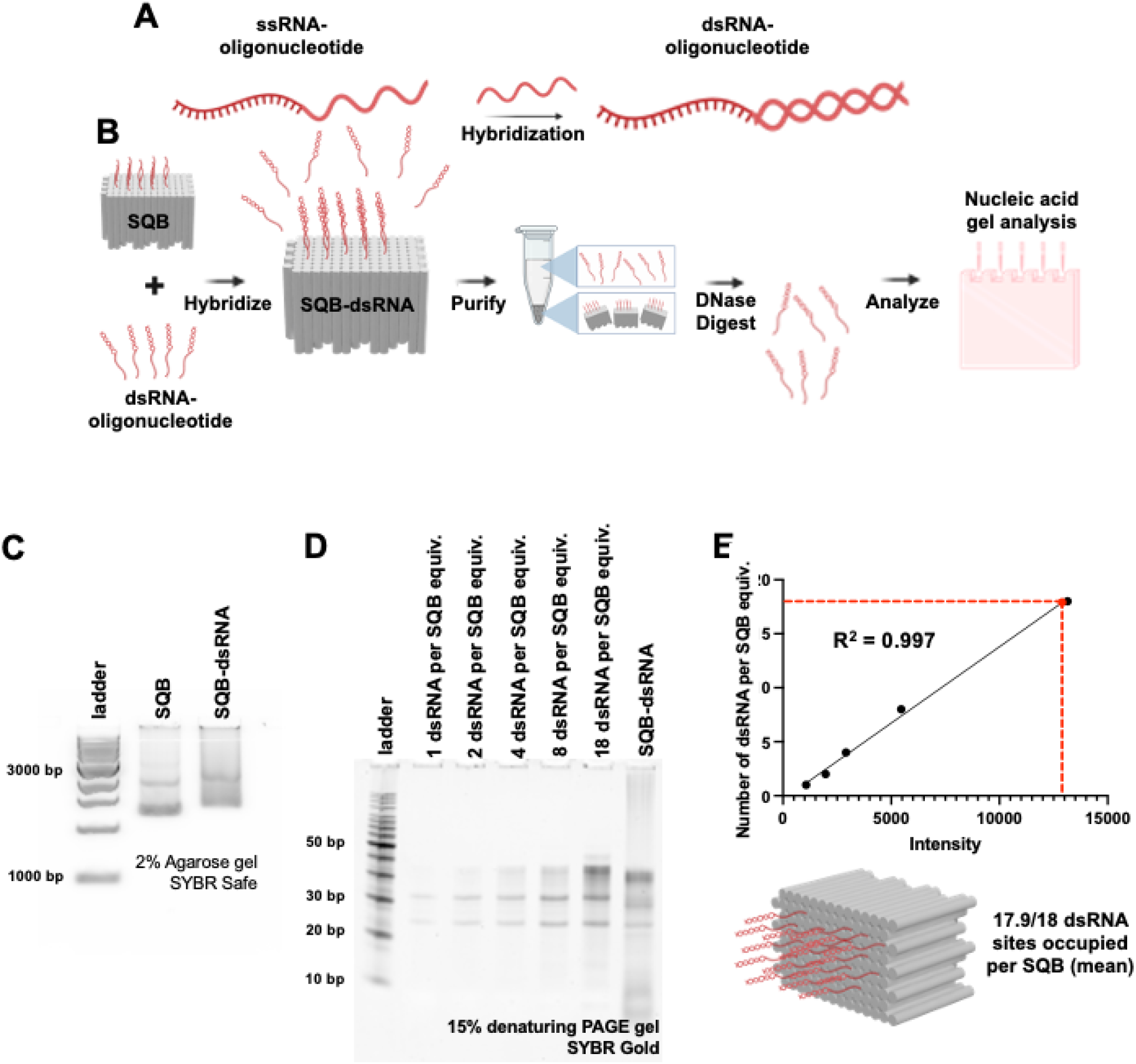
Quantifying double-stranded RNA (dsRNA) on DNA origami nanoparticles. (A) Schematic illustrating the design of the dsRNA for conjugation to the SQB, including one strand with an anti-handle DNA sequence complementary to a handle on the SQB. (B) Schematic illustrating the process of conjugating the dsRNA onto the SQB nanoparticle. (C) Agarose gel demonstrating the successful conjugation of dsRNA molecules on the SQB particle and the subsequent gel-based quantification analysis. (D) 15% denaturing PAGE gel of the SQB-dsRNA particles following DNase I digestion and post-staining with SYBR Gold; “equiv” refers to a unit of measure corresponding to the number of moles of SQB loaded in the rightmost lane, used as a reference. (E) Linear regression curve represents the relationship between the band intensity and the number of dsRNA molecules conjugated per SQB, as determined by ImageJ. This analysis quantifies the average occupancy of the 18 designed dsRNA-capture sites on the DNA origami samples, demonstrated for varying concentrations (1, 2, 4, 8, and 18 dsRNA per SQB).

### Conjugation and quantification of multiple independent cargos on DNA origami nanoparticles in parallel

We have successfully quantified diverse cargos, including peptides, proteins, and nucleic acids. Since therapeutic efficacy often benefits from co-delivery of multiple cargos on a single nanoparticle, we aimed to demonstrate the simultaneous quantification of DNA, RNA, and protein via a gel assay. We utilized different conjugation strategies for each cargo type: CpG oligonucleotides were directly modified on ‘staple’ strands, siRNA were hybridized using a ‘handle/anti-handle’ hybridization as described in the dsRNA quantification section, and OVA protein was conjugated with appropriate ‘anti-handle’ oligonucleotides using SMCC chemistry (Fig. 5A). Successful conjugation of each cargo types was confirmed by a band shift on agarose gel (Fig. 5B). The agarose gel demonstrated aggregation of the SQB after conjugation of the siRNA and the OVA protein. However, subsequent TEM imaging validated that the SQBs were monodispersed and well-formed, suggesting that the charge modifications due to conjugation of cargo affected gel motility (Supplementary Figure 8). To quantify the cargos, we performed a DNase I digestion assay to degrade the DNA components. This allowed us to visualize the modified CpG and siRNA through SYBR safe staining (Fig. 5C) and OVA protein via silver staining (Fig. 5D) on separate denaturing PAGE gels. Quantification using ImageJ revealed an average of 17.6 siRNA molecules per SQB out of 20 available sites (Fig. 5E, Supplementary Table 7, Supplementary Fig. 5), approximately 15.8 CpG oligonucleotides per SQB out of 18 sites (Fig. 5F, Supplementary Table 8, Supplementary Fig. 6), and about 12.2 OVA proteins per SQB out of the 24 available sites (Fig. 5G, Supplementary Table 9, Supplementary Fig. 7). These results confirm the feasibility of simultaneously quantifying multiple cargos on a single SQB nanoparticle using a DNase I assay coupled with ImageJ analysis of gel results. The occupancy rates of approximately 88% for both CpG and siRNA suggest that most sites are effectively occupied by the cargos, with full occupancy potentially limited by steric hindrance. The observed average of 12.2 OVA proteins per SQB aligns with previous data (Fig. 1-3), validating our earlier findings and suggesting that less than the maximum 24 sites are typically occupied by OVA protein due to steric constraints.

**Figure 5.**
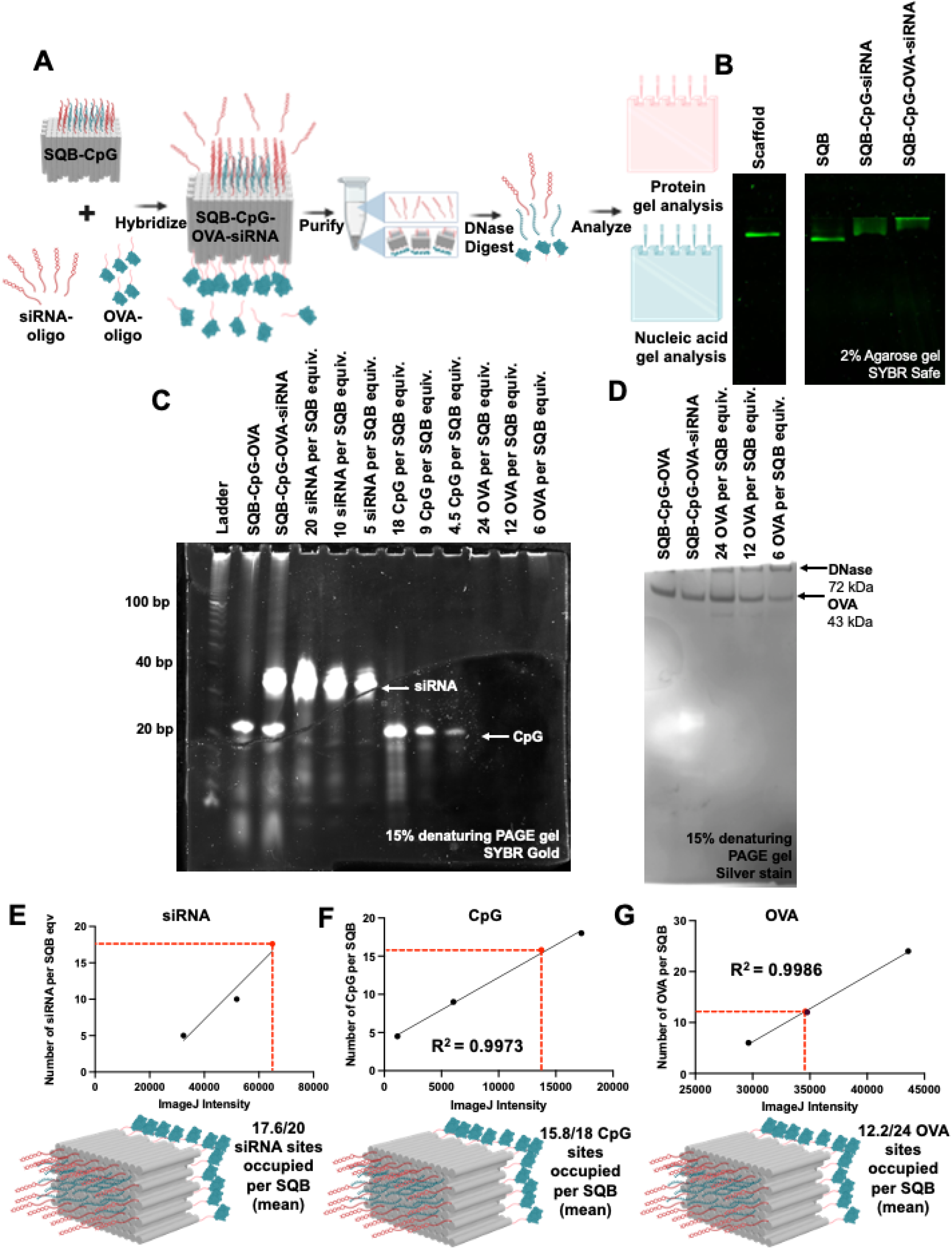
Quantifying cargo in a multiplexed fashion. (A) Schematic illustrating the conjugation of siRNA and OVA protein onto the CpG-SQB and the subsequent gel-based quantification analysis. (B) Agarose gel demonstrating the successful conjugation of siRNA and OVA molecules on the CpG-SQB nanoparticle. (C) 15% denaturing PAGE gel confirming the successful conjugation of siRNA and CpG oligonucleotides after DNase I degradation of the DNA origami scaffold and staple strands, followed by the analysis of the remaining cargo via SYBR Safe staining. Here, “equiv” refers to a unit of measure corresponding to the number of moles of SQB loaded in the left lane labeled SQB-CpG-OVA. The gel was cropped to highlight the bands of interest. (D) 15% denaturing PAGE gel confirming the successful conjugation of the OVA protein after DNase I degradation of the DNA origami scaffold and staple strands, with the remaining cargo analyzed via silver staining. “Equiv” refers to a unit of measure corresponding to the number of moles of SQB loaded in the leftmost lane. (E) Quantification of the siRNA on the DNA origami SQB, indicating5 and 10 siRNA molecules per SQB, via linear regression of the ImageJ band intensity and the number of occupied siRNA-capture sites. The resulting relationship between band intensity and the number of occupied siRNA-capture sites was used to determine the mean occupancy of the 20 designed siRNA-capture sites on the DNA origami samples. (F) Quantification of the CpG on the DNA origami SQB, indicating 4.5, 9, and 18 CpG molecules per SQB, via linear regression of the ImageJ band intensity and the number of occupied CpG-capture sites. This analysis provided the mean occupancy of the 18 designed CpG-capture sites on the DNA origami samples. (G) Quantification of the OVA protein on the DNA origami SQB, showing 6, 12 and 24 OVA molecules per SQB via linear regression of the ImageJ band intensity and the number of occupied OVA-capture sites. The resulting relationship between the band intensity and the number of occupied OVA-capture sites was used to determine the mean occupancy of the 24 designed OVA-capture sites on the DNA origami samples.

#### Concluding Remarks

With promising results from pre-clinical studies^13,16,34,40,41,53-55^, DNA origami appears well primed for advancing towards clinical translation as a precision therapeutic platform. However, the translation from laboratory research to clinical use, including achieving FDA approval, requires robust standards in chemistry, manufacturing, and controls (CMC) management. This process includes accurate methods for the fabrication, purification, and characterization of the therapeutic nanoparticle. A key advantage of DNA origami nanoparticles is their ability to conjugate a wide variety of cargos, including peptides, proteins, nucleic acids, small molecules, lipids, and carbohydrates. This versatility enhances their potential for therapeutic applications but also induces additional complexity, which poses a challenge for regulatory approval^56^. Therefore, precise quantification of the conjugated therapeutic cargos on DNA origami nanoparticles is crucial for their successful clinical translation.

In this study, we methodically describe two robust methods for quantifying virtually any cargo on DNA origami nanoparticles: gel-based and fluorescence-based methods. These methods are versatile and practical, allowing for the independent quantification of each type of cargo simultaneously on DNA origami nanoparticles. In summary, these quantification methods are crucial for advancing DNA origami towards clinical adoption while ensuring compliance with rigorous regulatory standards. They provide a framework for the accurate characterization of DNA origami nanoparticles, paving the way for their successful use in therapeutic applications.

## Acknowledgements

We appreciate the support and experimental input from many Shih Lab members. Special thanks to Kow Simpson, Kathleen Mulligan, and Michal Walzcak for their assistance with these studies, as well as Maurice Perez, and Michael Carr for their expertise with facilities and instrumentation. Figure schematics were generated using BioRender (https://biorender.com/).

## Author contributions

This work was written by Olivia J. Young*, Hawa Dembele*, Anjali Rajwar*, Yang C. Zeng, and William M. Shih. Y.C.Z. developed the idea for this manuscript. Y.C.Z., O.J.Y., H.D and A.R. planned experiments. O.J.Y., H.D and A.R. performed experiments. O.J.Y., H.D and A.R. drafted the manuscript with the guidance of Y.C.Z.

I.C.K. and J.H.R. provided guidance on the project and feedback on the manuscript.

W.M.S. and Y.C.Z. provided overall guidance for the project, and edited the manuscript.

## Data and materials availability

The paper and supplementary materials contain the supporting data for this study.

## Materials and Methods

### Material sources

Scaffold strand was harvested in-house from the M13 bacteriophage. All short oligonucleotide ‘staple’ strands, including strands modified with CpG or strands modified with an amine group for conjugation with various cargo, were purchased from IDT (www.idtdna.com). Peptides were purchased from Genscript with an azide-modified lysine on the N terminal. Proteins including ovalbumin (OVA) and OVA-AF488 were purchased from Sigma (#A2512) and ThermoFisher (O34781) respectively. GFP mRNA was purchased from TriLink Biotechnologies (L-7601-1000). The dsRNA and siRNA were obtained from IDT (www.idtdna.com) and then annealed into a double-stranded structure. Sequences for all cargos can be found in Supplementary Table 10.

### DNA origami nanoparticle fabrication

This study utilized the DNA origami square block (SQB), assembled from 126 double helices through complementary base pairing of 245 short staple oligonucleotides, purchased from IDT (www.idtdna.com), to the p8634 scaffold obtained from M13 bacteriophage^57^, following established protocols. The SQB was folded in buffer consisting of 5mM Tris, 1 mM ethylenediaminetetraacetic acid (EDTA; pH 8.0), 12 mM MgCl_2_ using an 18-hour thermocycler program, which denatured the strands at 80°C for 15 minutes, then annealing via a temperature ramp from 50°C to 40°C decreasing at -0.1°C every 10 minutes and 48 seconds. Detailed information is available in prior publications^12,13^. The 126 double helices have overhang ‘handle’ strands which enable plug-and-play conjugation of cargos onto the SQB. Cargo was conjugated onto a single-stranded ‘anti-handle’ oligonucleotide using a click chemistry strategy and then hybridized to the corresponding ‘handle’ strand via complementary base-pairing. After fabrication, DNA origami nanoparticles were purified from excess staple strands before conjugating with cargo as detailed below.

### Protein conjugation on DNA origami nanoparticles

Proteins were conjugated to an oligonucleotide via SMCC chemistry and then attached to the DNA origami nanoparticle via ‘handle/anti-handle’ base-pairing. An anti-handle oligonucleotide, synthesized with a 5’-NH2-TTCTAGGGTTAAAAGGGGACG-3’ sequence from IDT, was modified with a succinimidyl 4-(N-maleimidomethyl)cyclohexane-1-carboxylate (SMCC) linker, and then purified using a NAP column (GE Healthcare Life Sciences). The SMCC linker, equipped with NHS ester and maleimide groups, enables selective conjugation between amines and cysteines. The maleimide group forms a stable thioether bond by reacting with cysteine residues on the protein, while the NHS ester group forms an amide bond between the oligo and the SMCC linker by reacting with the oligo’s amine group. Unconjugated oligonucleotides were effectively removed by repeated washing steps using a 30K Amicon filter (Sigma). The conjugated OVA-oligo was then incubated with the DNA origami nanoparticle at 37°C for 2 hours, followed by removal of unbound OVA-oligo using PEG precipitation as described below.

### Peptide conjugation on DNA origami nanoparticle

Peptides were connected to DNA origami nanoparticles using a ‘handle’-’anti-handle’ approach, where the ‘handle’ oligo protruded from the DNA origami surface and the ‘anti-handle’ oligonucleotide, conjugated with cargo, hybridized via Watson-Crick base-pairing. Employing DBCO-Azide copper-free click chemistry, the ‘anti-handle’ oligonucleotide was linked to the peptide: the amine-modified oligonucleotide was reacted with Dibenzocyclooctyne-N-hydroxysuccinimidyl ester (DBCO-NHS ester; Millipore, #761524) in phosphate buffer pH 8.0, purified via Illustra NAP Column (GE Healthcare Life Sciences, #17-0852-02) and concentrated using 3K Amicron (Millipore; #UFC500324). The successful DBCO conjugation was verified through NanoDrop OD310 peak, and concentration was measured via NanoDrop. Peptide-azide (Gensript) was reacted overnight with the oligonucleotide-DBCO at a ratio of 1.5:1 as determined in prior experiments, and confirmed by a 15% denaturing PAGE gel (as described below). The peptide-oligonucleotide was purified using 8% PAGE gel extraction, filtering through Freeze ‘N Squeeze DNA Gel Extraction Spin Columns (Bio-Rad; #7326165) and then finally performing ethanol precipitation. Concentration of the peptide-oligonucleotide was calculated using the NanoDrop. Purity of the sample was determined by running a 15% dPAGE gel with the unpurified and purified sample. Peptide-oligo products can be stored in -20°C for several months. After purification, the peptide-oligo was hybridized onto DNA origami in 1XTE 10 mM MgCl2 buffer, incubated at 37°C for 1–2 hours, and then subjected to PEG precipitation. The resulting conjugated DNA origami underwent analysis via agarose gel electrophoresis, TEM, and DNase I assay for confirmation.

### Nucleic acid conjugation on DNA origami nanoparticle

#### CpG

CpG was incorporated into the DNA origami nanoparticles by extending the staple strands on the flat side of the DNA origami SQB. The CpG oligonucleotides were ordered with a phosphorothioate (PS) backbone (5’ – TCCATGACGTTCCTGACGTT-3’, IDT) for increased nuclease resistance at the 5’ end of the staple strands corresponding to designated positions in the precise spacing of 3.5 nm, determined previously to be optimal for immune stimulation^13^. These strands were incorporated into the DNA origami folding mixture as described above.

#### dsRNA and siRNA

dsRNA and siRNA were hybridized to the DNA origami nanoparticle via Watson-Crick base pairing. Prior to conjugation with RNA, the nanoparticle was assembled as described above. The RNA strands were ordered with a DNA-sense RNA hybrid and a short complementary anti-sense RNA strand from IDT. Then DNA-sense RNA hybrid contains a DNA region on the 3’ end and an RNA region on the 5’. The 5’ end of the double helices on the flat side of the SQB were extended to include staple strands complementary to the ssDNA region of the RNA duplex. RNA duplexing was performed by mixing both strands and incubating the mixture at 37°C for 30 min in PBS. For conjugation to the DNA origami nanoparticle, the RNA strands were added at a 4× excess and incubated on ice for 1 hour, then placed on a shaker at 37°C for 30 min.

### DNA origami nanoparticle purification

We utilized PEG precipitation for molecular weight-based separation of DNA origami nanoparticles from excess staple strands and cargos and concentration of the sample, both after DNA origami nanoparticle fabrication and after cargo conjugation. A 15% w/v PEG-8000 (Fisher Scientific, BP2331) buffer with 10 mM MgCl_2_ was prepared in 1× TE buffer. The PEG buffer was mixed in a 1:1 volume ratio with the nanoparticles, incubated for 30 minutes, and centrifuged at 16000g for 25 minutes, as described in a prior publication^13^. After discarding the supernatant, the pellet containing purified nanoparticles was resuspended in 1X TE buffer with 10 mM MgCl_2_ through shaking at 800 rpm for 30 minutes. Nanoparticle concentration was determined using NanoDrop, while sample purity was confirmed using agarose gel electrophoresis and TEM.

### DNase degradation

For the purpose of quantifying the cargo attached to the DNA origami, we relied on a DNase I assay to digest the DNA origami structure. We incubated 1 µg of DNA origami nanoparticle with 1 U/µL DNase I (2,000 units/mL, New England Biolabs, M0303S) with 10× DNase I buffer diluted in water (Gibco). In parallel, different amounts of the cargos matching different stoichiometries (still amounting to 1 µg of DNA origami sample) were prepared to undergo a DNase digestion as well. Samples were incubated at 37°C for 30 minutes to degrade the DNA.

### Gel confirmation

#### Agarose gel

Functionalized DNA origami samples were analyzed via 2% native agarose gel electrophoresis. The gel was prepared in 0.5× TBE with 11 mM MgCl_2_ and 0.005% v/v SYBR safe and run at 70V for 2 hours. If fluorescent cargo is functionalized on the DNA origami, SYBR Safe is not added, as the gel can be imaged directly without additional staining. Agarose gels were imaged with a Typhoon Gel Scanner and images analyzed via ImageJ.

#### Denaturing PAGE gel

Denaturing PAGE gels are used to analyze nucleic acid cargo after DNase I degradation of the DNA origami nanoparticle. Denaturing PAGE gel (15%) was made in-house using 9 mL urea concentrate (Fisher Scientific), 4.5 mL urea dilutant, 1.5 mL urea buffer, 10 µL tetramethylethylenediamine (TEMED) and 150 µL 10 wt% ammonium persulfate (APS) cast into a mini cassette (ThermoFisher Novex). The percent urea in the gel can be adjusted depending on the application. 1–5 picomoles of samples (after DNase degradation) were mixed with formamide loading buffer (FLB). The samples were denatured at 70°C for 10 min, the samples were loaded and the gel was run in 0.5× TBE buffer for approximately 45 min at 250 V. The gel was removed from the cassette, stained with SYBR Gold for 5–10 minutes and then imaged with the Sapphire Gel Scanner. Subsequent quantification was performed using ImageJ as described below.

#### SDS-PAGE gel

SDS-PAGE gel is used to analyze oligonucleotide conjugation to protein and also to analyze protein and peptide cargo after DNase I degradation of the DNA origami nanoparticle. Once the samples were prepared for analysis, they were mixed with 4× NuPAGE LDS sample buffer (ThermoFisher; #NP0008), incubated at 95°C for 2 minutes in the thermocycler, and loaded in 4–12% NuPAGE Bis-Tris gels (ThermoFisher; #NP0322). An SDS-PAGE gel was run in 1× MES SDS running buffer (ThermoFisher; #NP0002) at 150 V for 45 minutes. If a fluorescent tag such as AF488 was conjugated to the protein, the gel could be imaged directly on the Sapphire gel imager under the SYBR safe setting, as AF488 has approximately the same wavelength as SYBR. For non-fluorescent proteins and peptides, the gel was subjected to silver stain (Pierce, #24612), following the manufacturer’s protocol. Imaging was performed using the Silver Stain setting on Image Gel 6 on Gel Doc EZ Imager (Bio-Rad). ImageJ software was used for band intensity quantification as described below.

### TEM imaging

The DNA origami nanoparticles’ morphology and structure were examined using negative stain transmission electron microscopy (TEM), providing high-resolution imaging for comprehensive analysis. To prepare samples, carbon-coated grids underwent a 30-second plasma discharge treatment to ensure surface cleanliness. A 3– 4 µL solution containing DNA origami nanoparticles (4–10 nM) was applied to the grid and allowed to adsorb for 45 seconds. Excess solution was removed via gentle blotting with filter paper, followed by addition and removal of a 2–3 µl of 0.75% uranyl formate solution to enhance contrast. A second drop of uranyl formate was applied for a two-minute incubation period and then wicked away using filter paper. The grid was them applied for TEM analysis, utilizing a JEOL 1400 TEM operating in brightfield mode at 120 kV. TEM enabled visualization and detailed characterization of nanoparticle structural features, including size, shape, distribution, and potential aggregation or abnormalities.

### Fluorescence quantification of DNA origami nanoparticles

To quantify OVA protein conjugation, a standard curve was established by correlating AF488 fluorescence intensity (measured via Nanodrop) with the molar concentration of AF488-labeled OVA (ThermoFisher, O34781). Iterative experiments systematically varied OVA concentrations, correlating AF488-labeled OVA fluorescent intensities to known OVA molar concentrations. Using linear regression in GraphPad Prism, we were able to derive a reliable equation comparing OVA concentration with AF488 intensity measurements, which was used to calculate subsequent OVA molar concentrations accurately. Using the established equation, AF488 intensity was used to estimate the OVA content on the DNA origami nanoparticles. Dividing this concentration by the DNA origami nanoparticle concentration (determined via NanoDrop) yielded an approximate count of OVA molecules per nanoparticle, offering insights into the final product’s composition. OVA loading efficiency was determined by comparing the number of OVA on each DNA origami nanoparticle to the theoretical number of sites for OVA conjugation.

### Gel quantification of conjugated cargo via Image J

ImageJ was employed for gel electrophoresis band quantification. The gel bands were selected using the rectangular tool (Ctrl+1), then shifted across lanes using Ctrl+2. Plotting lanes and their corresponding bands was accomplished by choosing Analyze->Gels->Plot lanes or using Ctrl+3, generating a plot for each lane. To minimize background noise, we drew a baseline with the line tool and connected peaks to the baseline. We selected each peak using the wand tracing tool. The selected peaks’ areas were calculated sequentially and displayed in the result window. We determined the conjugation efficiency by dividing the total area of conjugated protein (DNA origami particle with protein) by the area of unconjugated protein (protein only), yielding an estimation of conjugation efficiency. The detailed image J analysis and quantification can be found in the supplementary materials.

### Statistical analysis

GraphPad Prism 10 was used to run fitting analysis and graphics presentation. To correlate the gel intensity results with the theoretical number of occupied cargo-capture sites on the origami, we used the ‘simple linear regression’ function to generate a best-fit line and then to run inference on unknown quantities.

