## Supplementary Information for "Cargo quantification of functionalized DNA origami for therapeutic application"

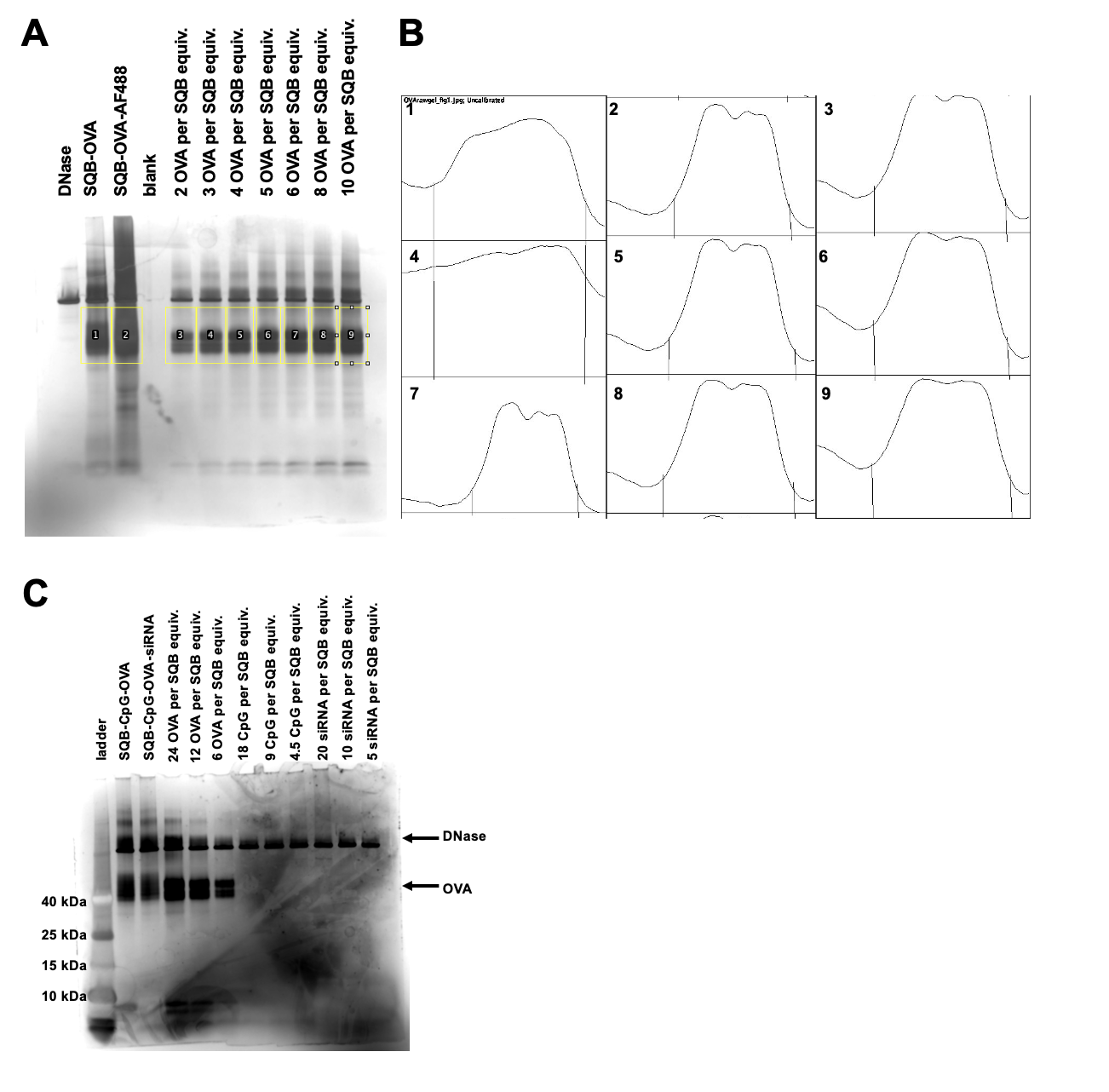
**Supplementary Figure 1 | Raw data and band intensity peaks associated with Figure 1F, as measured by ImageJ. (A)** Raw gel with labeled lanes. **(B)** ImageJ band intensity peaks. This intensity measurement of OVA in the SDS-PAGE gel allowed us to compare it with the theoretical intensity of 4, 5, 6 and 8 OVA molecules (n=1 gel image analysed) via the equation: *Number of OVA per SQB = 0.0008085 x Intensity – 31.45*. **(C)** Raw gel of the protein ladder. Ladder was aligned, relying on the DNase band for alignment with the gel of interest.

**B**

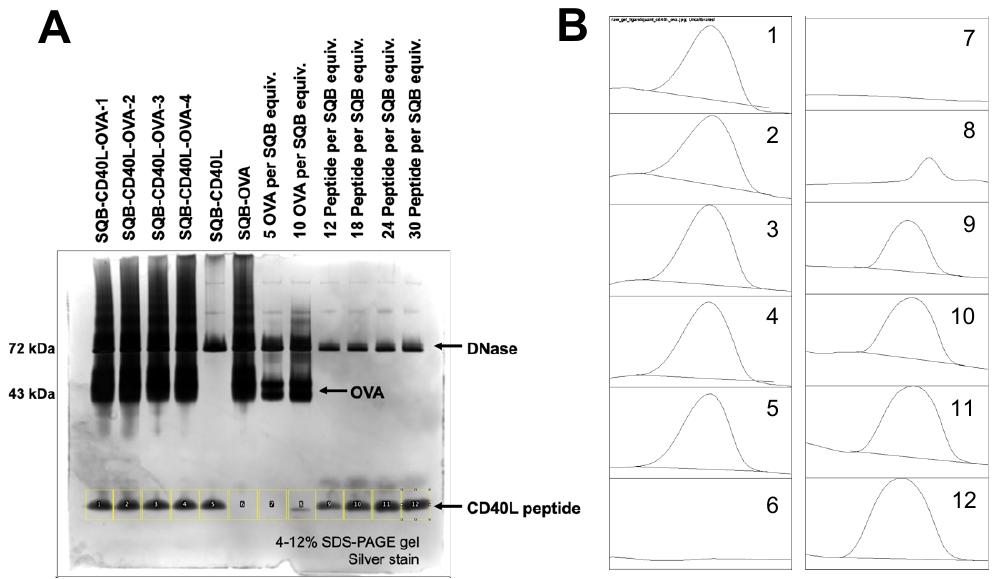

**Supplementary Figure 2| Raw data and band intensity peaks associated with Figure 3F, as measured by ImageJ. (A)** Raw gel with labeled lanes. **(B)** Band intensity peaks. This intensity measurement of peptide in the SDS-PAGE gel allowed us to compare it with the theoretical intensity of 18 peptide molecules (n=1 gel image analyzed) via the equation: *Number of CD40L peptide per SQB = 0.000792 x Intensity – 1.806.*

**A**

**B**

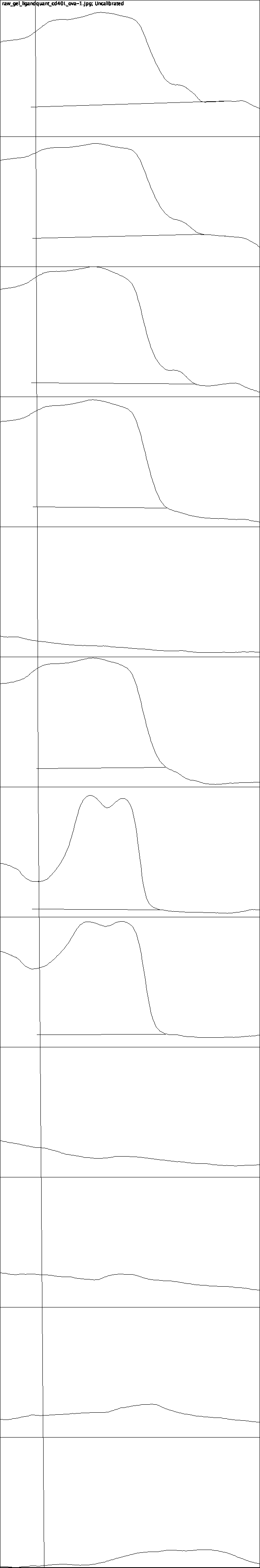

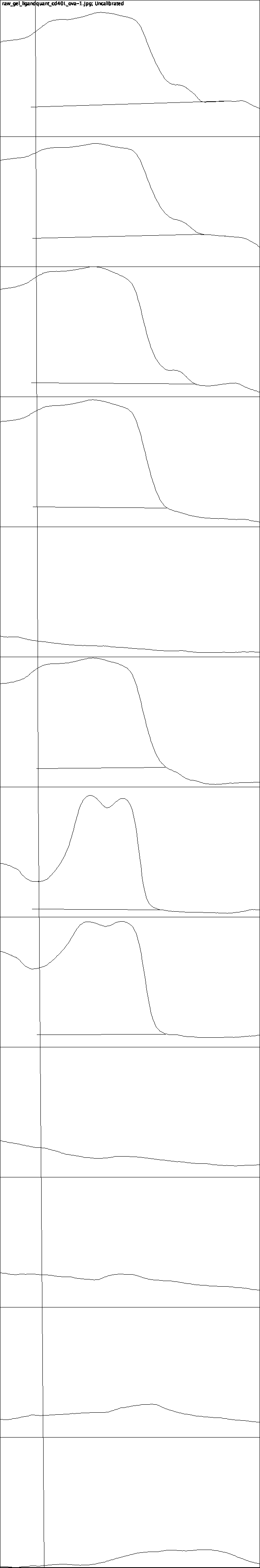

1
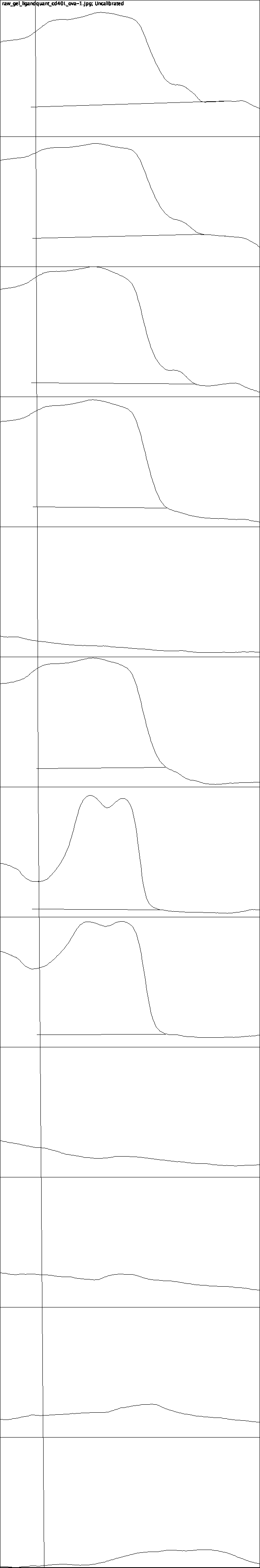

2
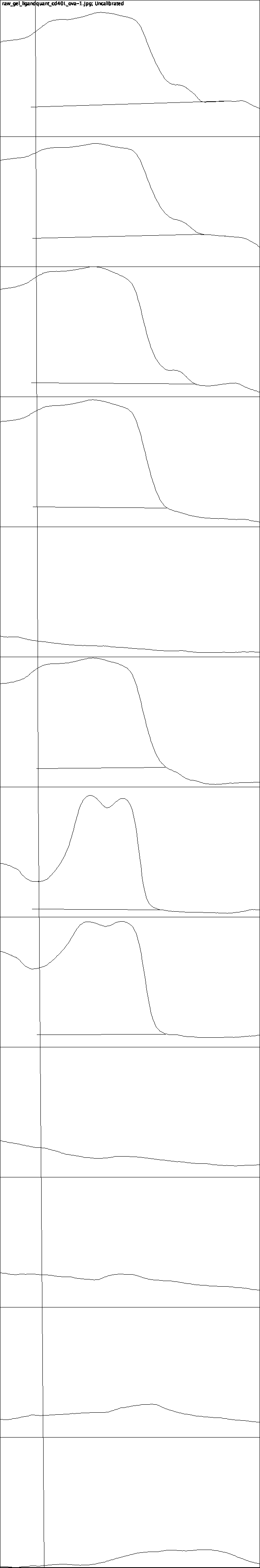

3
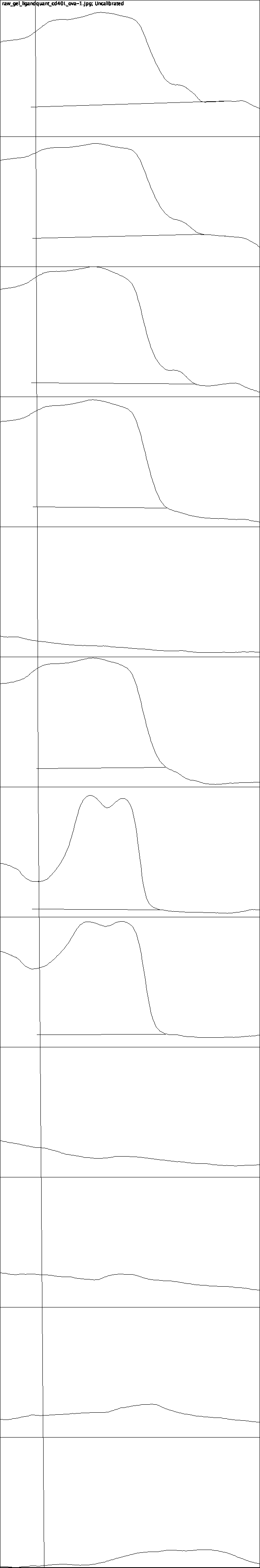

4
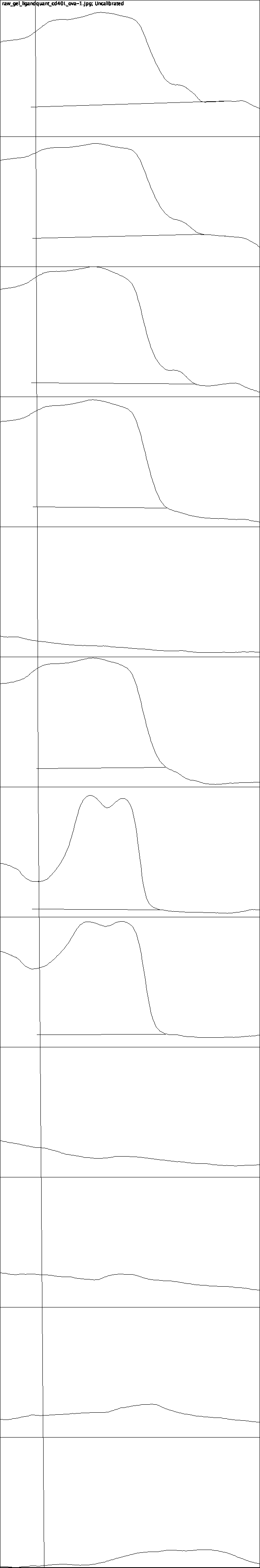

5
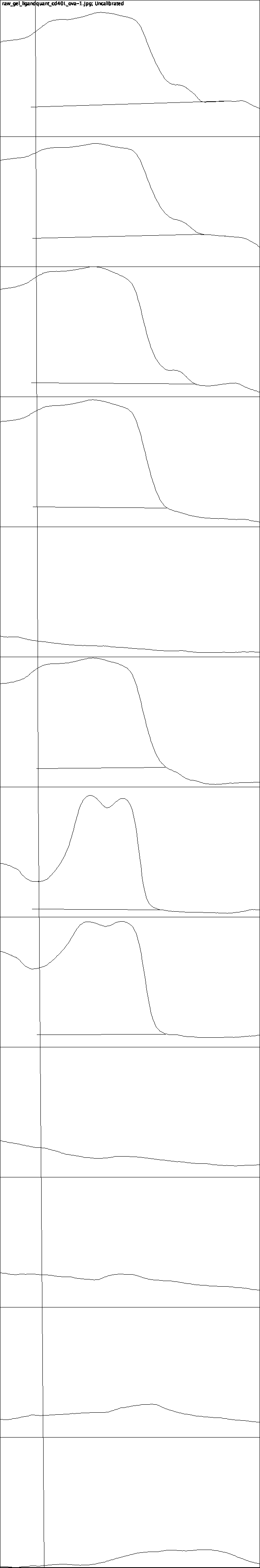

6
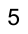

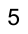

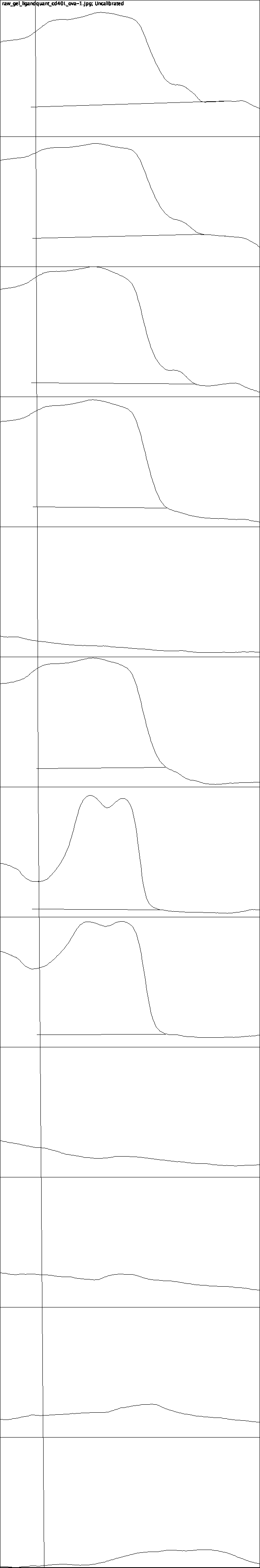

7
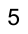

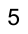

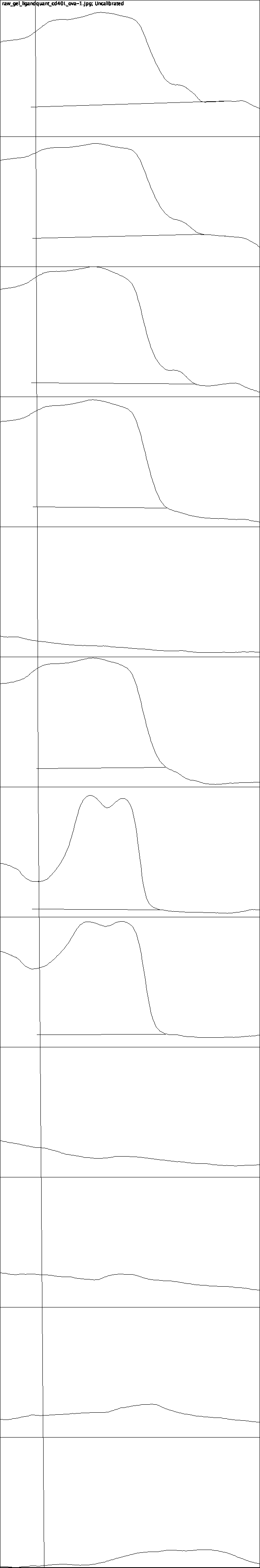

8
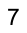

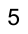

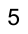

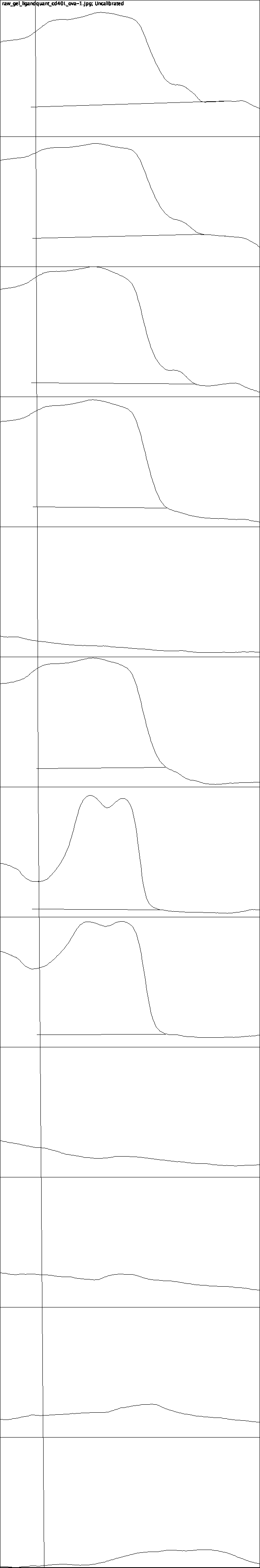

9
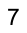

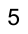

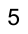

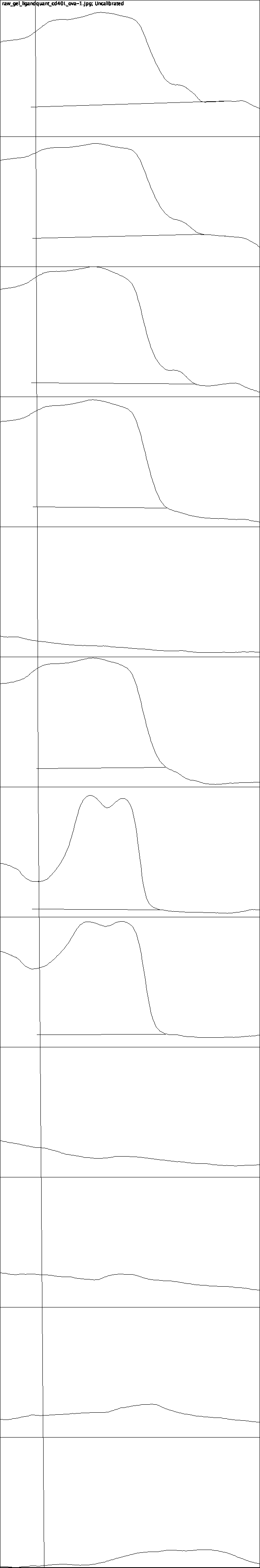

10
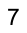

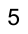

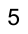

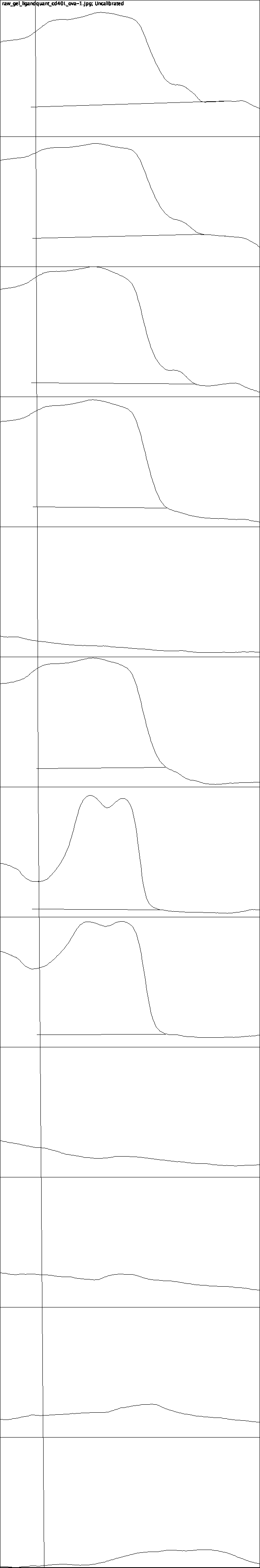

11
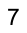

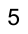

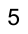

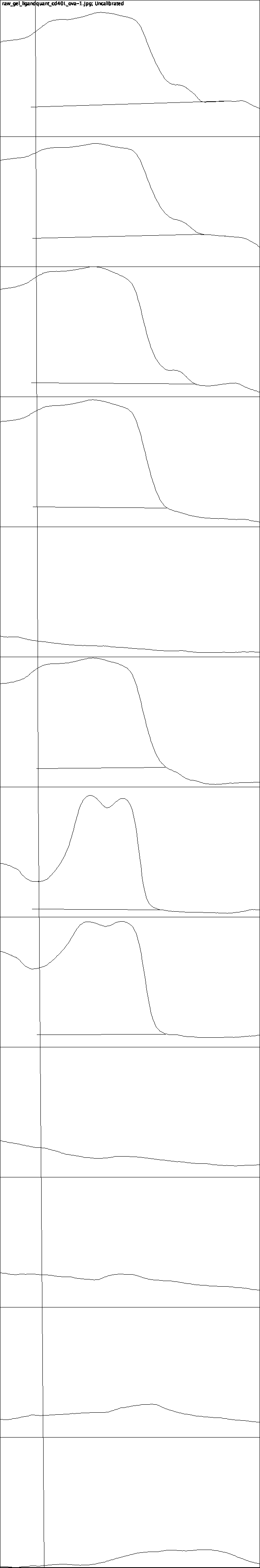

12
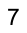

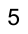

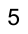

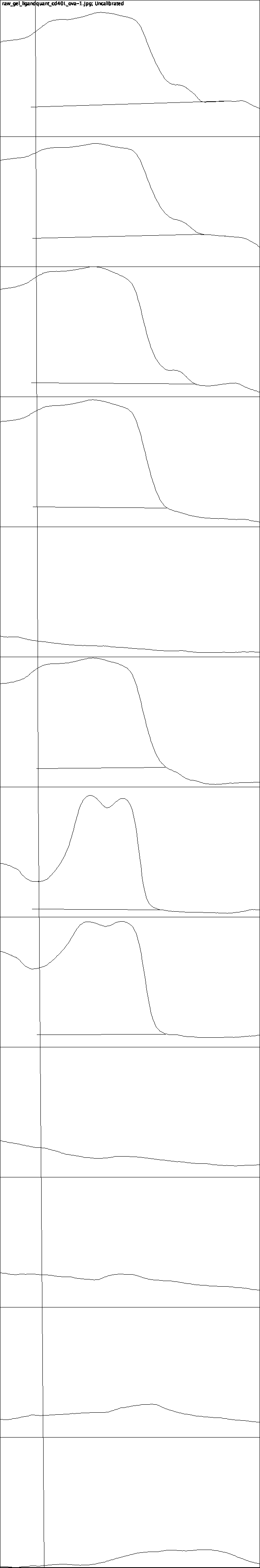

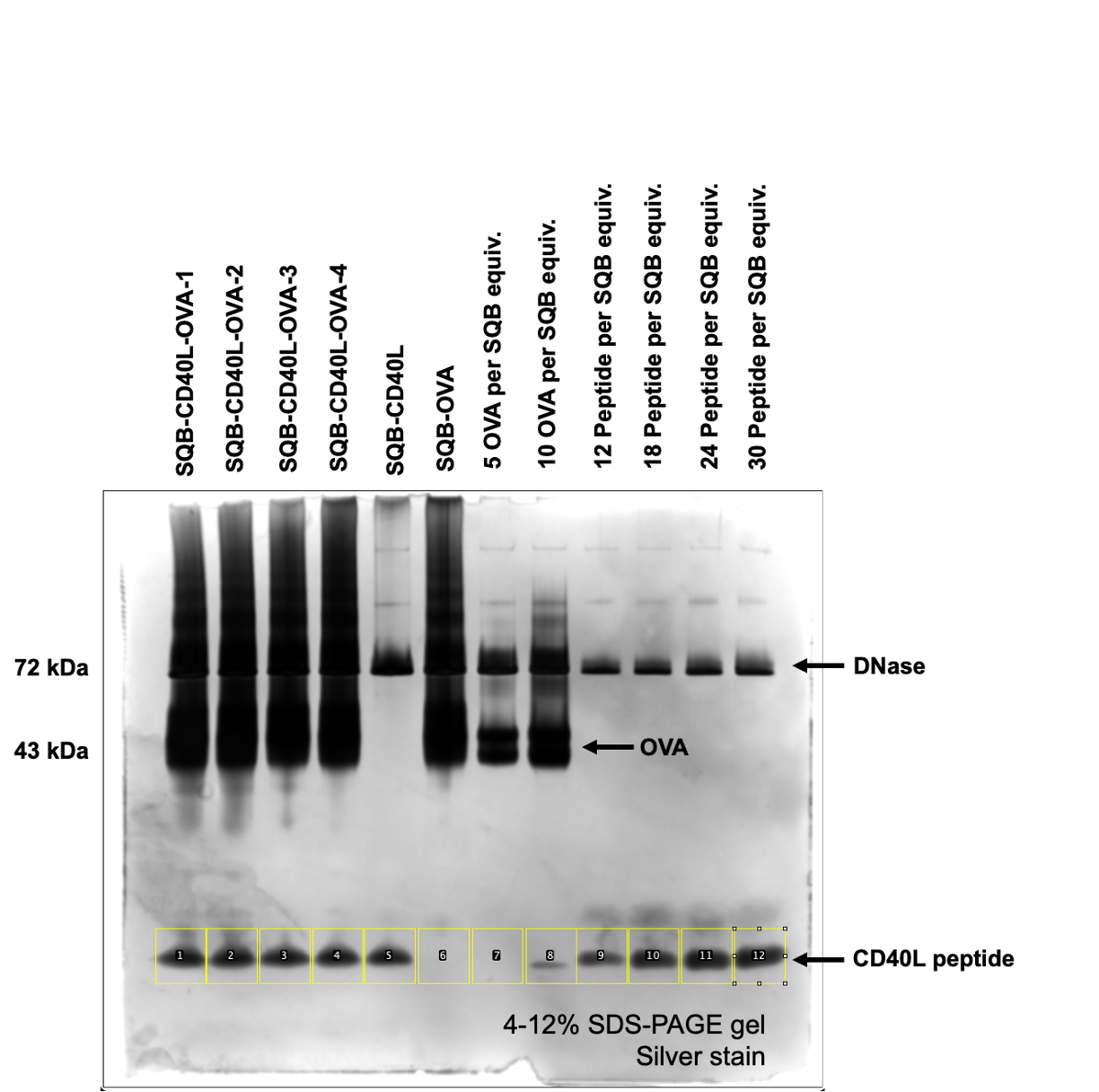

**C**

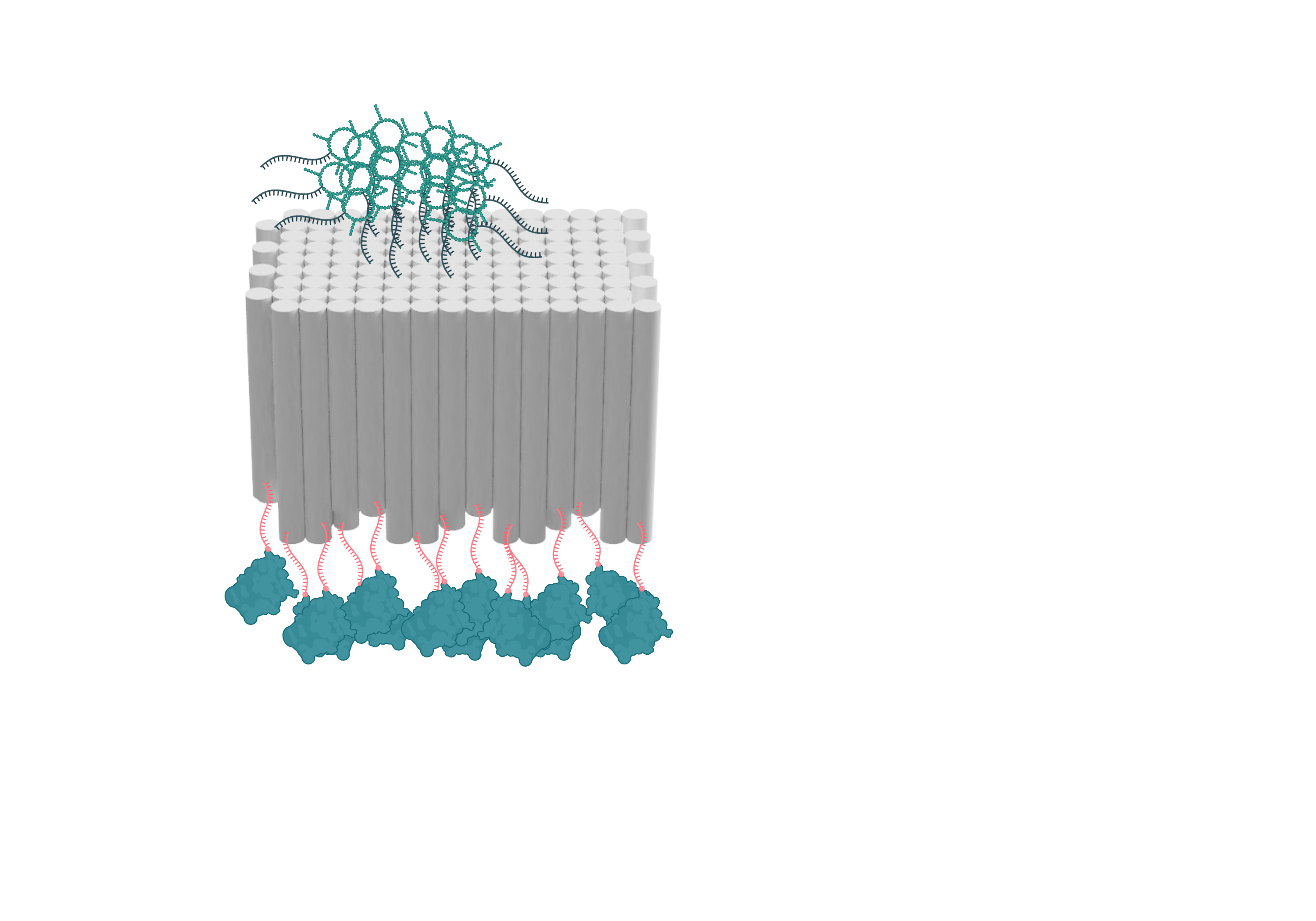

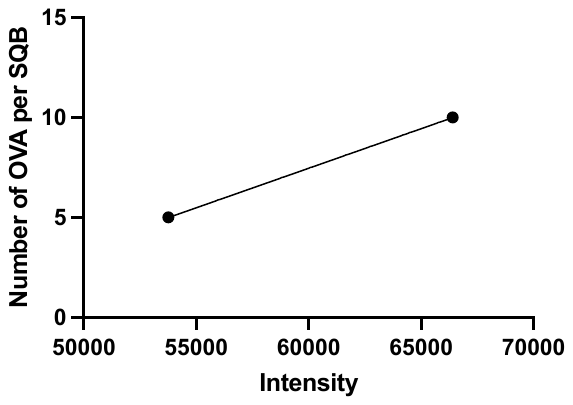

**Number of OVA per SQB =**

**0.0003955 x Intensity – 16.26**

**11.8 OVA**

**per SQB**

**Supplementary Figure 3| Band intensity peaks associated for OVA quantification associated with Figure 3 as measured by ImageJ. (A)** Raw gel with labeled lanes. **(B)** Band intensity peaks. This intensity measurement of OVA in the SDS-PAGE gel allowed us to compare it with the theoretical intensity of 24 OVA molecules (n=1 gel image analyzed). **(C)** Quantification of the OVA protein on the DNA origami SQB relying on linear regression of the ImageJ band intensity and the theoretical percent of conjugated ligand sites on the DNA origami. The resulting relationship between the band intensity and the percent of conjugated sites was used to determine the percent of conjugated sites on the DNA origami samples.

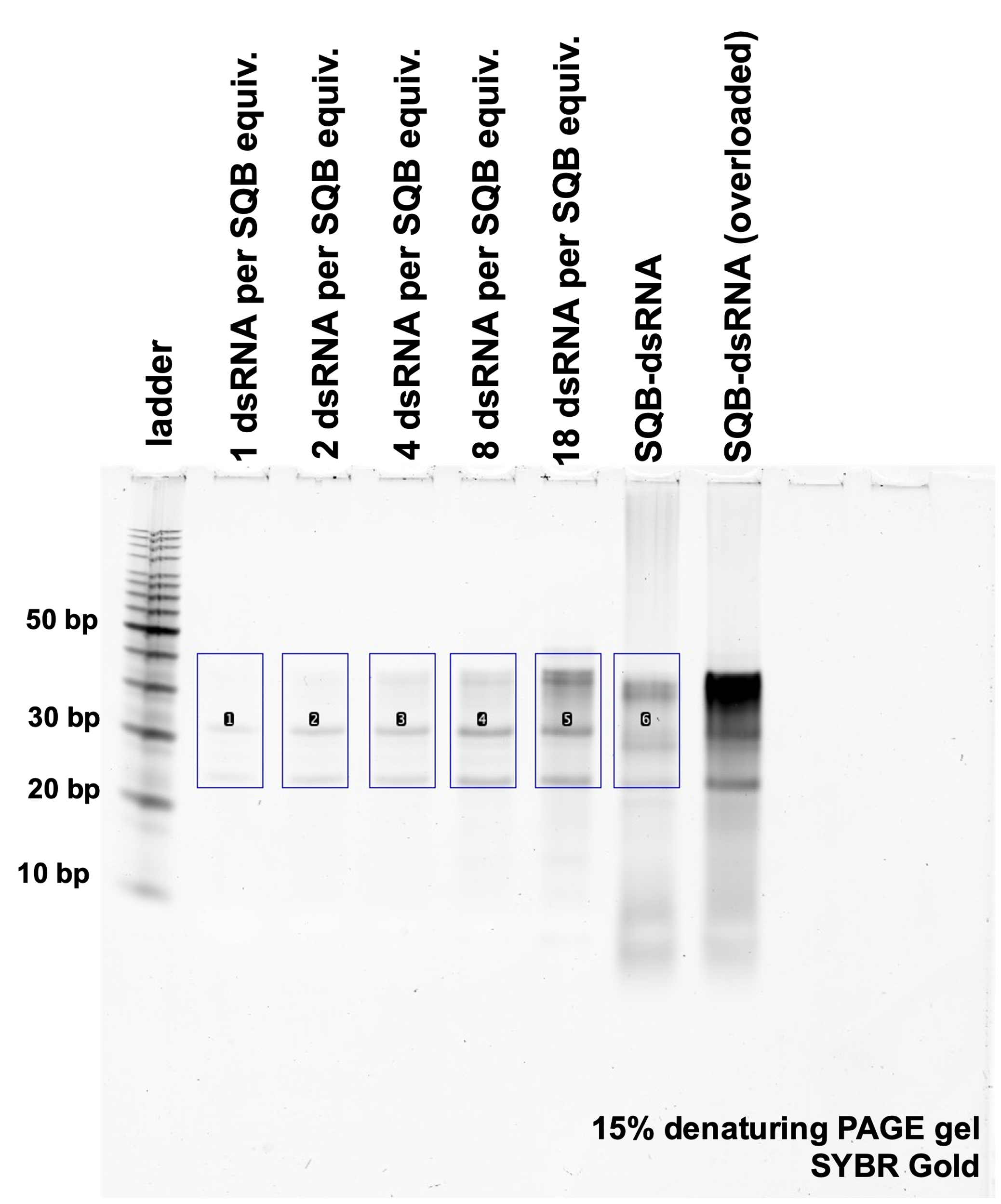

**A**

**B**

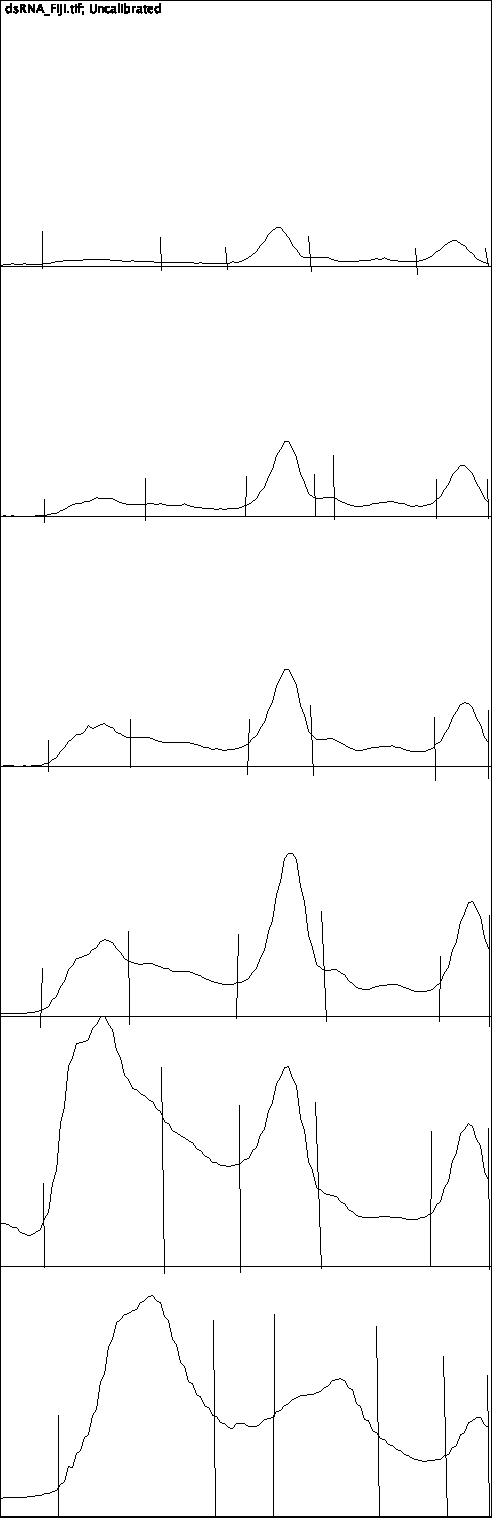

1 dsRNA per SQB

2 dsRNA per SQB

4 dsRNA per SQB

8 dsRNA per SQB

12 dsRNA per SQB

SQB-dsRNA

**Supplementary Figure 4| Band intensity peaks associated with Figure 4E, as measured by ImageJ. (A)** Raw gel with labeled lanes. **(B)** Band intensity peaks. This intensity measurement of dsRNA in the SDS-PAGE gel allowed us to compare it with the theoretical intensity of dsRNA molecules (n=1 gel image analyzed) via the equation: *Number of dsRNA per SQB = 705.68 x Intensity – 256.07*.

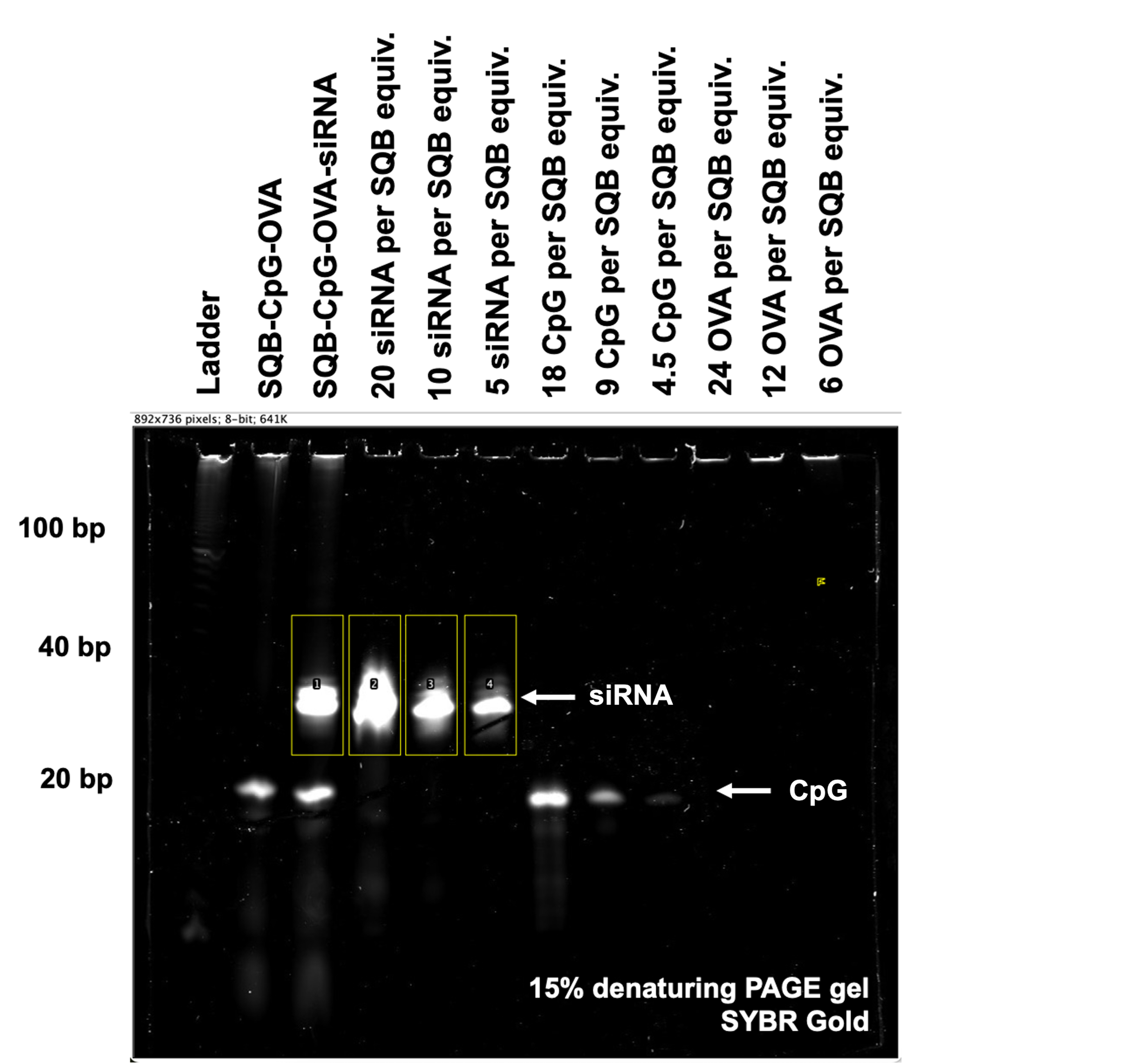

**A**

**B**

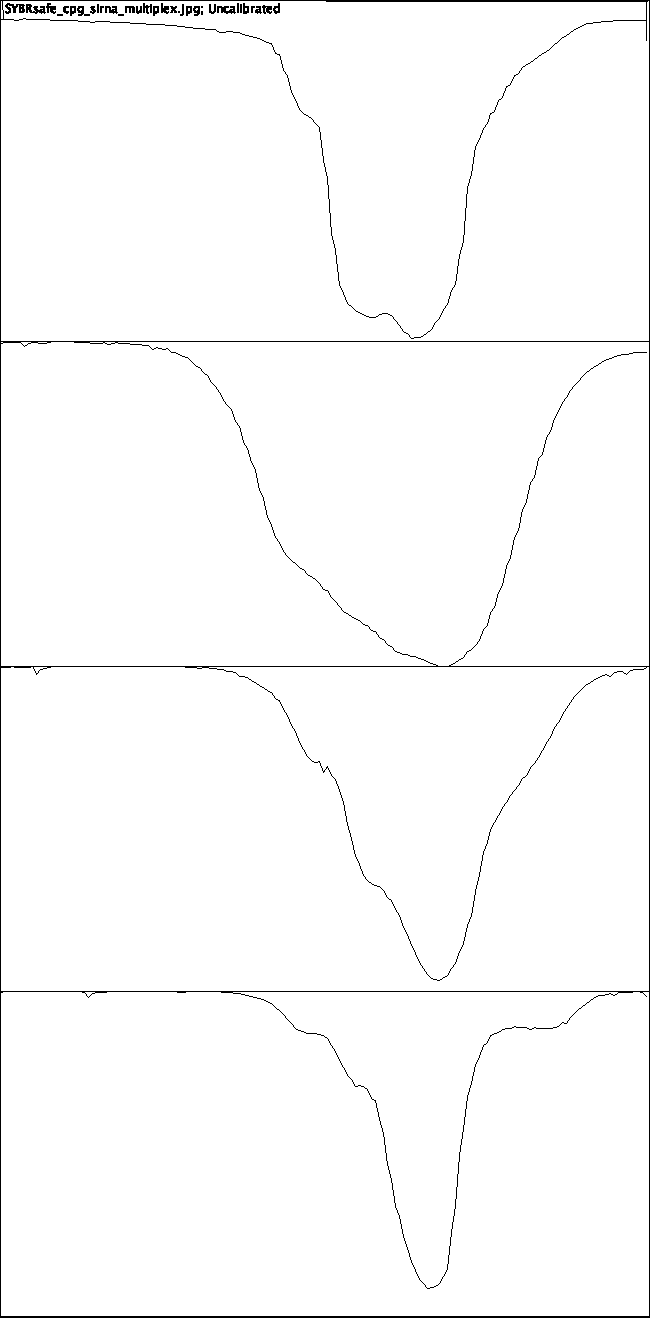

SQB-CpG-OVA-siRNA

20 siRNA per SQB

10 siRNA per SQB

5 siRNA per SQB

**Supplementary Figure 5| Band intensity peaks associated with Figure 5D, as measured by ImageJ. (A)** Raw gel with labeled lanes. **(B)** Band intensity peaks. This intensity measurement of siRNA in the SDS-PAGE gel allowed us to compare it with the theoretical intensity of 20 siRNA molecules (n=1 gel image analyzed) via the equation: *Number of siRNA per SQB = 0.0002565 x Intensity - 3.304*. Due to the smeared nature of the 20 siRNA per SQB band, this band was excluded from the subsequent linear regression analysis.

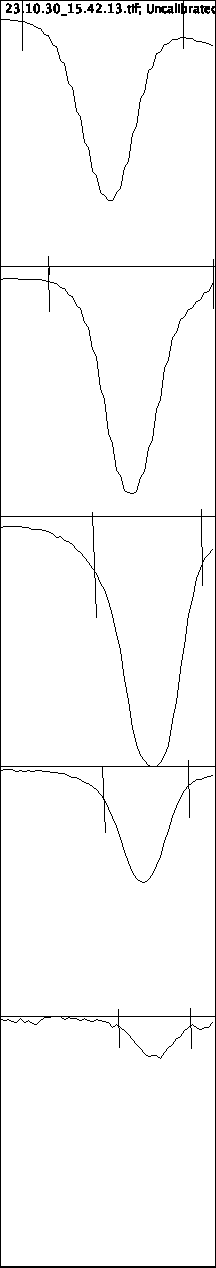

**B**

**A**

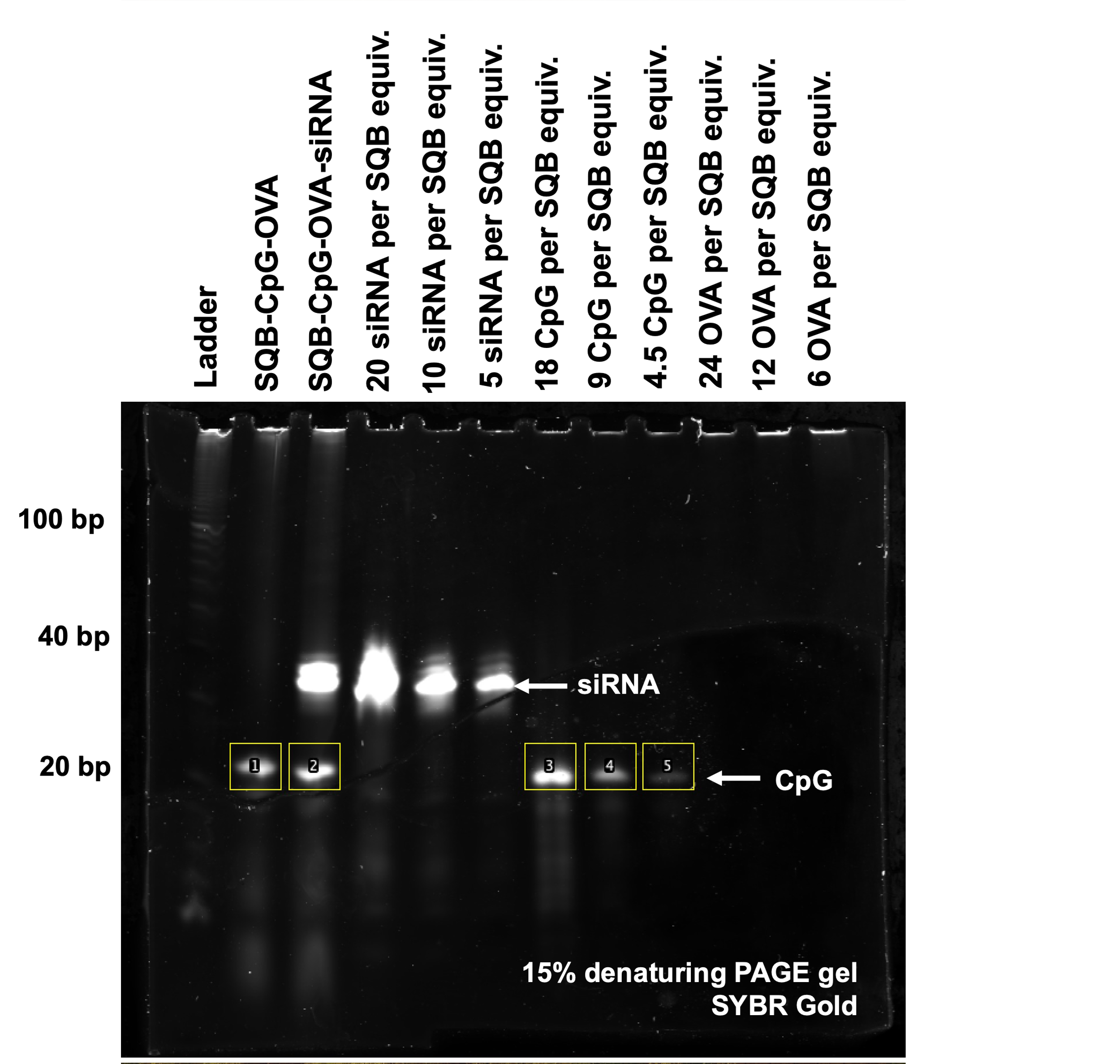

SQB-CpG-OVA

SQB-CpG-OVA-siRNA

18 CpG

9 CpG

4.5 CpG

**Supplementary Figure 6| Band intensity peaks associated with Figure 5E, as measured by ImageJ. (A)** Raw gel with labeled lanes. **(B)** Band intensity peaks. This intensity measurement of CpG in the denaturing PAGE gel allowed us to compare it with the theoretical intensity of 18 CpG molecules (n=1 gel image analysed) via the equation: *Number of CpG per SQB = 0.0008654 x Intensity + 3.097*.

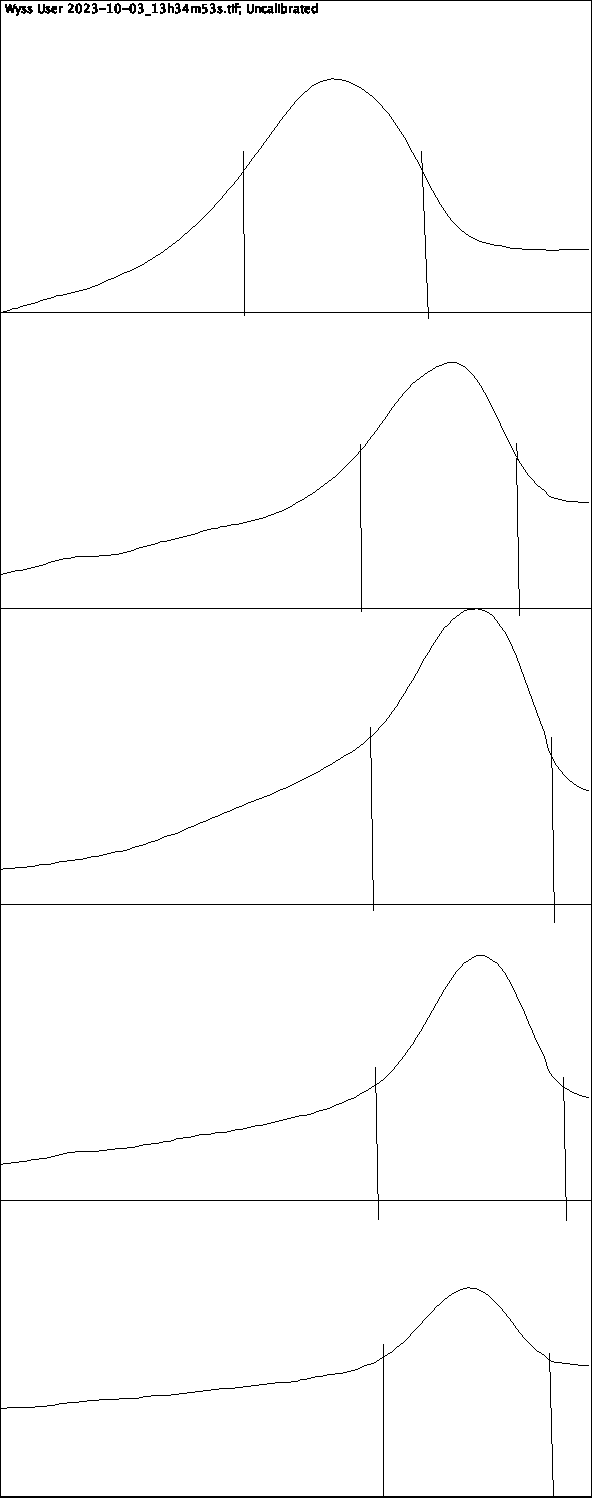

**A**

**B**

SQB-CpG-OVA

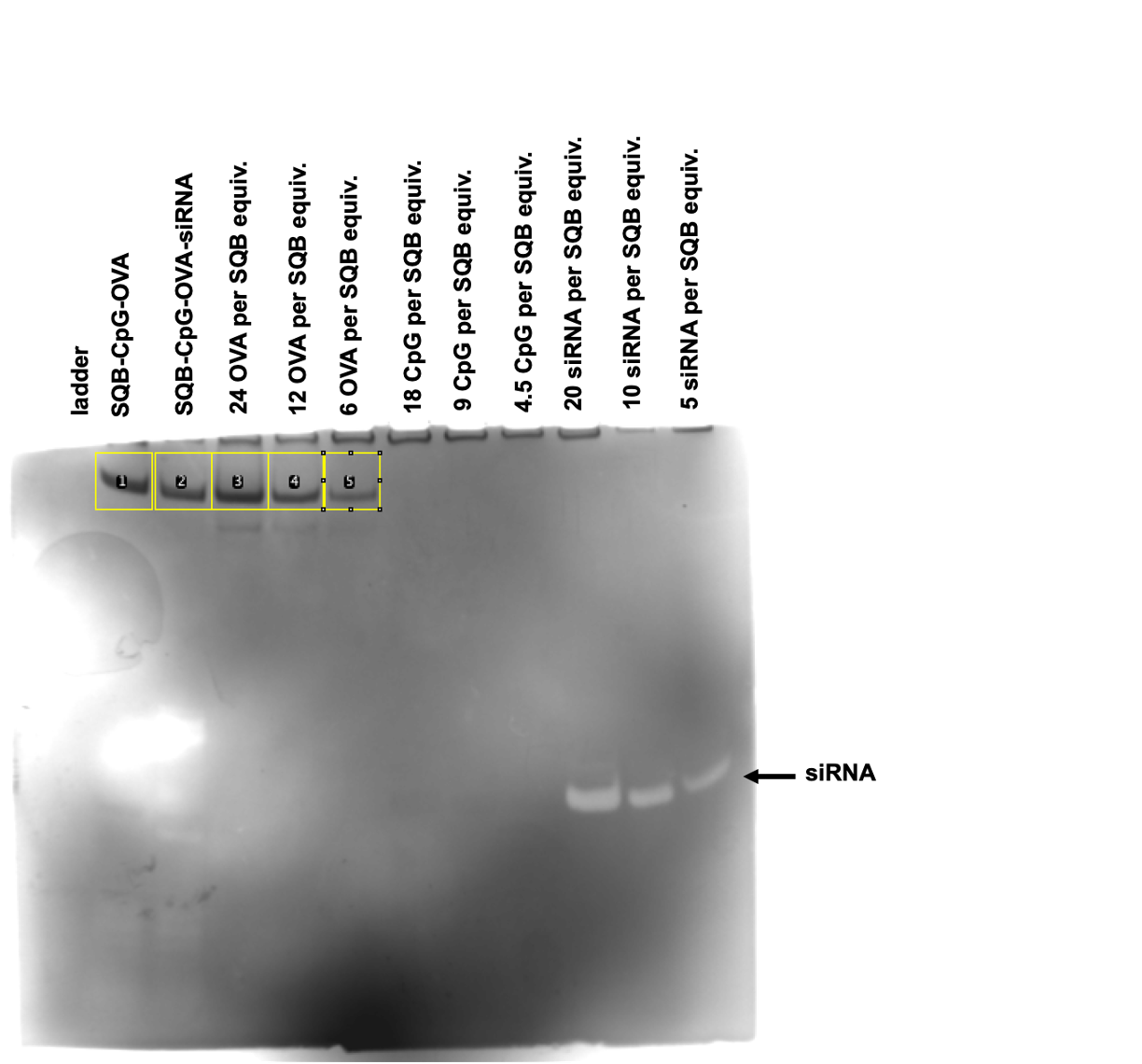

SQB-CpG-OVA-siRNA

24 OVA per SQB

12 OVA per SQB
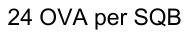

6 OVA per SQB
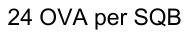

**Supplementary Figure 7| Band intensity peaks associated with Figure 5F, as measured by ImageJ. (A)** Raw gel with labeled lanes. **(B)** Band intensity peaks. This intensity measurement of OVA in the SDS-PAGE gel allowed us to compare it with the theoretical intensity of 24 OVA molecules (n=1 gel image analysed) via the equation: *Number of OVA per SQB = 0.001299 x Intensity – 32.76*.

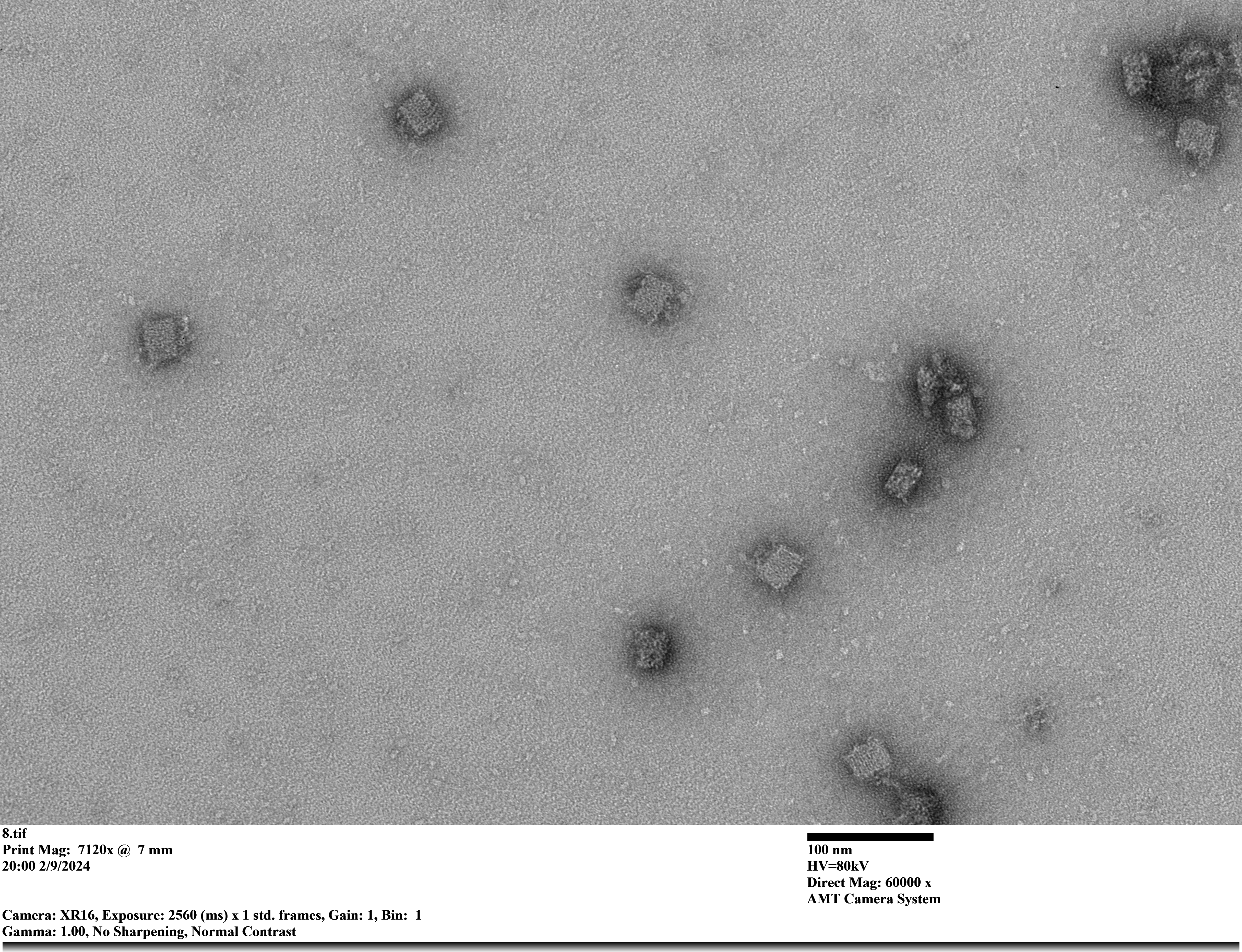

**Supplementary Figure 8| TEM image of the SQB conjugated to OVA, SiRNA, and CpG.** TEM imaging revealed mostly monodispersed SQB nanoparticles. Images were acquired at a magnification of 60000x.

| **Lane** | **ImageJ Intensity** | **Number of OVA per SQB** |
| --- | --- | --- |
| 4 OVA per SQB equiv. | 43853.158 |  |
| 5 OVA per SQB equiv. | 44982.158 |  |
| 6 OVA per SQB equiv. | 46491.38 |  |
| 8 OVA per SQB equiv. | 48730.622 |  |
| SQB-OVA | 53469.451 | 11.7800511 |
| SQB-OVA-AF488 | 64976.054* | 21.0831397* |
|  | **Average** | 11.8 |

**Supplementary Table 1 | Raw data for Figure 1E, comparing the theoretical number of OVA per SQB to the OVA-conjugated DNA origami nanoparticle via SDS-PAGE analysis.** Intensity analyzed by ImageJ analysis. The SQB-OVA lane has an amount of OVA protein equivalent to 164.5 femtomoles, which in this case is referred to as an “equiv”. *SQB-OVA-AF488 could not be accurately quantified due to the high presence of background in this lane, likely caused by the AF488 fluorophore, so this value was excluded from the analysis.

| **Structure** | **260/280** | **ng/uL** | **MW** | **uM** | **nM** | **Number of OVA per SQB equiv.** | **Optimal density of AF488 (pmol/ul)** |
| --- | --- | --- | --- | --- | --- | --- | --- |
| Oligo-OVA (8.8 uM) | 1.72 | 43.6 | 6629.31 | 8.8 | 8800 | 101.1 | 52.8 |
| Oligo-OVA (4.4 uM) | 1.73 | 21.6 | 6629.31 | 4.4 | 4400 | 50.6 | 27.1 |
| Oligo-OVA (2.2 uM) | 1.74 | 9.6 | 6629.31 | 2.2 | 2200 | 25.3 | 12.8 |
| Oligo-OVA (1.1 uM) | 1.7 | 3.1 | 6629.31 | 1.1 | 1100 | 12.6 | 4.7 |
| Oligo-OVA (0.55 uM) | 1.7 | 2.1 | 6629.31 | 0.55 | 550 | 6.3 | 3.6 |
| Oligo-OVA (0.275 uM) | 1.69 | 0.6 | 6629.31 | 0.275 | 275 | 3.2 | 2.1 |

**Supplementary Table 2 | Raw data for Figure 2E, comparing the theoretical concentration of OVA to the optical density of the AF488 fluorescence from OVA-AF488.** The AF488 fluorescence optical density (OD) was measured via the microarray setting on the Nanodrop. We assumed a SQB concentration of 87 nM, averaged from the SQB concentrations tested in Supplementary Table 3, to calculate the number of OVA per SQB equivalent.

| **Structure** | **A260/**  **A280** | **ng/uL** | **nM** | **AF488 (pmol/ul)** | **OVA uM** | **OVA nM** | **SQB nM** | **OVA/SQB** |
| --- | --- | --- | --- | --- | --- | --- | --- | --- |
| SQB-OVA-AF488 | 1.7 | 869 | 159.43 | 12.9 | 2.18112 | 2181.12 | 159.43 | 13.68 |
| SQB-OVA-AF488 (1:2 dilution) | 1.69 | 506.9 | 93 | 6.6 | 1.14288 | 1142.88 | 93 | 12.29 |
| SQB-OVA-AF488 (1:4 dilution) | 1.69 | 276.6 | 50.75 | 3.6 | 0.64848 | 648.48 | 50.75 | 12.78 |
| SQB-OVA-AF488 (1:8 dilution) | 1.69 | 244.8 | 44.91 | 1.6 | 0.31888 | 318.88 | 44.91 | 7.1* |
| Average |  |  |  |  |  |  |  | 12.9 |

**Supplementary Table 3 |Calculations for Figure 2E, determining the number of OVA per SQB using the determined linear relationship between AF488 fluorescence and SQB concentration.** The AF488 fluorescence optical density (OD) was measured via the microarray setting on the Nanodrop, while the SQB concentration (in nM) was measured using the dsDNA setting on the Nanodrop. The determined equation is: *OVA concentration (uM) = 0.1648 x OD-AF488 + 0.05520*. *We did not include the 1:8 dilution data in the final average, as the dilution might have been more dilute than intended, leading to a vastly different number than the previous three data points.

| **Lane** | **ImageJ Intensity** | **Calculated number of CD40L peptide per SQB** |
| --- | --- | --- |
| SQB-CD40L-OVA-1 | 30296.53 | 22.2 |
| SQB-CD40L-OVA-2 | 30930.45 | 22.7 |
| SQB-CD40L-OVA-3 | 33632.53 | 24.8 |
| SQB-CD40L-OVA-4 | 32880.7 | 24.2 |
| SQB-CD40L | 31832.16 | 23.4 |
| CD40L 12 per SQB | 17260.2 |  |
| CD40L 18 per SQB | 25052.78 |  |
| CD40L 24 per SQB | 33007.04 |  |
| CD40L 30 per SQB | 39824.75 |  |
|  | **Average** | **23.5** |

**Supplementary Table 4|Raw data for Figure 3F, comparing the theoretical number of peptides per SQB to the peptide-conjugated DNA origami nanoparticle via SDS-PAGE analysis.** Intensity analyzed by ImageJ analysis.

| **Lane** | **ImageJ Intensity** | **Calculated number of OVA per SQB** |
| --- | --- | --- |
| SQB-CD40L-OVA-1 | 67665.149 | 10.5 |
| SQB-CD40L-OVA-2 | 67889.229 | 10.6 |
| SQB-CD40L-OVA-3 | 78770.593 | 14.9 |
| SQB-CD40L-OVA-4 | 70378.773 | 11.6 |
| SQB-OVA | 70347.187 | 11.6 |
| OVA 5 per SQB | 53759.999 |  |
| OVA 10 per SQB | 66402.614 |  |
|  | **Average** | **11.8** |

**Supplementary Table 5|Raw data for OVA analysis related to Supplementary Figure 3, comparing the theoretical number of OVA per SQB to the OVA-conjugated DNA origami nanoparticle via SDS-PAGE analysis.** Intensity analyzed by ImageJ analysis.

| **Number of dsRNA** | **ImageJ Intensity** | **Calculated number of dsRNA per SQB** |
| --- | --- | --- |
| 1 | 1069.422 |  |
| 2 | 1971.372 |  |
| 4 | 2913.247 |  |
| 8 | 5463.888 |  |
| 18 | 13149.828 |  |
| SQB-dsRNA | 12899.282 | 17.92 |
|  | **Average** | **17.92** |

**Supplementary Table 6 |Raw data for Figure 4E, comparing the theoretical number of dsRNA per SQB to the dsRNA-conjugated DNA origami nanoparticle via denaturing PAGE analysis.** Intensity analyzed by ImageJ analysis.

| **Lane** | **ImageJ Intensity** | **Number of siRNA per SQB** |
| --- | --- | --- |
| SQB-CpG-OVA-siRNA | 64947.011 | 17.6 |
| 10 siRNA | 51867.487 |  |
| 5 siRNA | 32375.191 |  |
|  | **Average** | **17.6** |

**Supplementary Table 7 |Raw data for Figure 5D, comparing the theoretical number of siRNA per SQB to the siRNA-conjugated DNA origami nanoparticle via 15% denaturing PAGE gel analysis.** Intensity analyzed by ImageJ analysis.

| **Lane** | **ImageJ Intensity** | **Number of siRNA per SQB** |
| --- | --- | --- |
| SQB-CpG-OVA | 15337.409 | 16.4 |
| SQB-CpG-OVA-siRNA | 16525.359 | 15.3 |
| 18 CpG | 17351.953 |  |
| 9 CpG | 6360.69 |  |
| 4.5 CpG | 1949.255 |  |
|  | **Average** | **15.8** |

**Supplementary Table 8 |Raw data for Figure 5E, comparing the theoretical number of CpG per SQB to the CpG-conjugated DNA origami nanoparticle via 15% denaturing PAGE gel analysis.** Intensity analyzed by ImageJ analysis.

| **Lane** | **ImageJ Intensity** | **Number of siRNA per SQB** |
| --- | --- | --- |
| SQB-CpG-OVA | 36131.48 | 14.2 |
| SQB-CpG-OVA-siRNA | 33089.874 | 10.2 |
| 24 OVA per SQB | 43580.087 |  |
| 12 OVA per SQB | 34761.158 |  |
| 6 OVA per SQB | 29642.409 |  |
|  | **Average** | **12.2** |

**Supplementary Table 9 |Raw data for Figure 5F, comparing the theoretical number of OVA per SQB to the OVA-conjugated DNA origami nanoparticle via SDS-PAGE analysis.** Intensity analyzed by ImageJ analysis.

| **Cargo type** | **Source** | **Sequence** |
| --- | --- | --- |
| Ovalbumin (OVA) | Sigma (#A2512) | MGSIGAASMEFCFDVFKELKVHHANENIFYCPIAIMSALAMVYLGAKDSTRTQINKVVRFDKLPGFGDSIEAQCGTSVNVHSSLRDILNQITKPNDVYSFSLASRLYAEERYPILPEYLQCVKELYRGGLEPINFQTAADQARELINSWVESQTNGIIRNVLQPSSVDSQTAMVLVNAIVFKGLWEKAFKDEDTQAMPFRVTEQESKPVQMMYQIGLFRVASMASEKMKILELPFASGTMSMLVLLPDEVSGLEQLESIINFEKLTEWTSSNVMEERKIKVYLPRMKMEEKYNLTSVLMAMGITDVFSSSANLSGISSAESLKISQAVHAAHAEINEAGREVVGSAEAGVDAASVSEEFRADHPFLFCIKHIATNAVLFFGRCVSP |
| CD40L cyclic peptide | Genscript custom order | {d-Lys(N3)}{Lys({Ahx}YYGK)}{D-ALA}{LYS({Ahx}YYGK)}{D-ALA}{LYS({Ahx}YYGK)} |
| dsRNA | IDT custom order | DNA handle+anti-sense RNA: AGTGATGTGAGACCATGTGAGrArCrCrArArGrGrCrArArUrArGrGrArArGrArArCrCrArUrCrU  Sense RNA: rArGrArUrGrGrUrUrCrUrUrCrCrUrArUrUrGrCrCrUrUrGrGrU |
| siRNA | IDT custom order | Sense: UGACCAAUUCAGCUGUAUG  Anti-Sense: CAUACAGCUGAAUUGGUCA |
| CpG | IDT custom order | TCCATGACGTTCCTGACGTT |

**Supplementary Table 10 |Source and sequence information for therapeutic cargos.**
